# Evolutionary analysis of LP3 gene family in conifers: an ASR homolog

**DOI:** 10.1101/2020.03.27.011197

**Authors:** Jonathan Lecoy, MR García-Gil

**Affiliations:** Department of Forest Genetics and Plant Physiology, Umeå Plant Science Center, SLU, Umeå, Sweden

**Author notes:** Corresponding author: Maria Rosario García-Gil.

**Keywords:** ASR, ABA/WDS, LP3, drought resistance, pine, selective pressures, codon usage, GC-content

## Abstract

Drought has long been established as a major environmental stress for plants which have in turn developed several coping strategies, ranging from physiological to molecular mechanisms. *LP3*; a homolog of the Abscisic Acid, Stress and Ripening (*ASR*) gene was first detected in tomato; and has been shown to be present in four different isoforms in loblolly pine called *LP3-0, LP3-1, LP3-2* and *LP3-3*. While *ASR* has already been extensively studied notably in tomato, the same cannot be said of *LP3*. Like *ASR*, the different *LP3* isoforms have been shown to be upregulated in response to water deficit stress and to also act as transcription factors for genes likely involved in hexose transport. In this study we have investigated the evolutionary history of *LP3* gene family, with the aim of relating it to that of *ASR* from a phylogenetic perspective and comparing the differences in selective pressure and codon usage. Phylogenetic analyses of different *LP3* homologs compared to *ASR* show that *LP3* is less divergent across species than *ASR* and that even when comparing the different sub-sections of the gene the divergence rate of *LP3* is lower than that of *ASR*. Analysis of different gene parameters showed that there were differences in GC1% and GC2% but not in total or GC3% content. All genes had a relatively high CAI value associated with a low to moderate ENC value, which is indicative of high translation efficiency found in highly expressed genes. Analysis of codon usage also showed that *LP3* preferentially uses different codons than *ASR*. Selective pressure analysis across most of the *LP3* and *ASR* genes used in this study showed that these genes were principally undergoing purifying selection, with the exception of *LP3*-*3* which seems to be undergoing diversifying selection most probably due to the fact that it likely recently diverged from *LP3*-*0*. This study thus provides insight in how *ASR* and *LP3* have diverged from each other while remaining homologous.

## 1. Introduction

Land colonisation by plants during the Paleozoic has forced these to adopt several adaptive strategies to survive desiccation (Edwards and Selden, 1992). These strategies led to the development of organs such as roots for taking up water and the implementation of water stress management tactics like the closure of stomata and the modulation of osmotic pressures within the plant cell in an effort to maintain the plants’ water potential (Chaves et al., 2003). Today, many plants species have adapted to be able to cope with drought through millennia of evolution, yet anthropogenic climate change is expected to dramatically affect the growth conditions of most plant species, notably through increased drought occurrence and aridity around the world.

Drought is a major hazard to the survival and development of commercially important plants, from both crops to forest tree species. In recent years, there has been an observed increase in drought occurrences notably in southern Europe, sub-Saharan Africa and many other areas around the world and this trend will only increase with time as climate change continues to progress (Gudmundsson and Seneviratne, 2016; Ruosteenoja et al., 2018). In this context, it is more important than ever to understand more about the mechanisms by which plants adapt and overcome water deficit stress in an effort to potentially produce more drought resistant varieties.

Water deficiency as a major stress for plant species is detected in many ways, with the signalling component being mediated largely through the phytohormone Abscisic acid (ABA) which is involved in stress response in plants, notably via its’ effects on gene expression and osmotic pressure adjustment within the plant cell (Bray, 1993). ABA is also implicated in the plant response to cold stress and the ripening process. The ABA dependent pathway has been the focus of extensive studies in a multitude of species, notably *Arabidopsis thaliana* L. and *Populus tremula* L.

Since the 1990s several research projects have focused on a drought responsive gene called ABA, Stress and Ripening (*ASR*), first detected in tomato leaves yet remarkably absent from *Arabidopsis thaliana* (Iusem et al., 1993). This research has led to the discovery of many different *ASR* orthologues and paralogs, with tomato having five different *ASR* genes, and rice up to six to date (Dominguez and Carrari, 2015; Frankel et al., 2006). *ASR*1 in tomato has by far been the most studied *ASR* gene in tomato. Transgenic expression of the *ASR* gene in Arabidopsis produced a phenotype similar to what is observed in *abi4* mutants in addition to an increased tolerance to salt, cold and other stresses (González and Iusem, 2014; Yang et al., 2005).

The *ASR* gene family contains a highly conserved midsection gene domain called the ABA/WDS domain (Pfam reference: PF02496), which is also highly conserved in *LP3*. The ABA/WDS domain is also expressed in mushrooms of the *Fomitopsis* genus, a membrane protein of *Pseudomonas* and angiomotin found in fern (Wang et al., 2002; Padmanabhan et al., 1997; González and Iusem, 2014). *ASR* in its native state is a disorganised, highly hydrophilic protein that requires two zinc ions to bind to lysine located in its N-terminal region to adopt its functional conformation, which leads to a protein dimerization and in turn bind to the plants’ DNA sequence (Goldgur et al., 2007; González and Iusem, 2014). *ASR* proteins act as transcription factors that induce the expression of aquaporines, cellulose synthases (CESA) and glucanases. *ASR*1, the most studied of the *ASR* genes, is for example involved in sugar metabolism in response to drought (Dominguez and Carrari, 2015).

An *ASR* homolog called *LP3* was first discovered in the roots of loblolly pine (*Pinus taeda* L.) whose background constitutive expression was significantly upregulated in drought conditions (González and Iusem, 2014; Padmanabhan et al., 1997; Wang et al., 2002). For the purposes of this study, while both genes are homologous, the terms *LP*3 and *ASR* will be used to describe the gymnosperm and angiosperm sequences, respectively. *LP*3 differs from *ASR* in that it contains a consequent insertion of 35 amino acids between its N-terminal and ABA/WDS regions and is present as a gene family (each individual gene isoform is called *LP*3-0, *LP*3-1, *LP*3-2 and *LP*3-3; *LP*3-2 and *LP*3-3 have only been partially sequenced therefore only partial sequences are available) within pines and other gymnosperms (Chang et al., 1996; Padmanabhan et al., 1997). *LP*3 transport into the nucleus is mediated by the putative C-terminal Nuclear Localisation Signal (NLS) of sequence KKESKEEEKEAEGKKHHH (Padmanabhan et al., 1997; Wang et al., 2002). Alternatively, this NLS sequence might not be necessary due to the short size of the *LP*3 protein which should allow it to diffuse through the nuclear envelope. Indeed research into *AS*R has shown that the NLS sequence to the one described above is not necessary for the *ASR* protein to diffuse into the nucleus (Ricardi et al., 2012). *LP*3 has not been as extensively studied as *ASR*, probably due to the difficulty of genetic studies within gymnosperms. This lack of study is the motivation for this study.

The objective of this research work is to investigate and compare the rate and mode of evolution of two orthologous genes *LP*3 and *ASR* genes. To achieve our objective we have conducted the following actions: We have (i) retraced the phylogeny of *LP*3 as a member of the ABA/WDS family and relate it to the *ASR* genes, (ii) estimated the GC content and Codon Usage Bias (CUB), and (iii) determined the mode of evolution of different subsections of the *ASR*/*LP*3 genes.

## 2. Materials and Methods

### 2.1 Identification of LP3 and ASR genes

Loblolly pine *LP*3-0, *LP*3-1, *LP*3-2 and *LP*3-3 CDSs were downloaded from NCBI *(Padmanabhan et al., 1997)*. Homologous angiosperm *ASR* and where possible gymnosperm *LP*3 whole gene CDSs were then extracted from NCBI using these sequences as queries via BLASTN in the NCBI database *(Boratyn et al., 2013)* with an e-value of 1^e-10^ as a threshold. More complete gymnosperm homologous sequences were also extracted via BLASTN in the Gymno Plaza database(v1.0) *(Altschul, 1997)*. For naming sequences in the instances where the gymnosperm sequences were uncharacterized, comparison with loblolly pine *LP*3 sequences via the NEEDLE alignment tool were done and whichever alignment had the highest score was used to determine to which isoform the uncharacterized sequence was most likely to be orthologous. Those sequences were then number as *LP*3-0-1, *LP*3-0-2, *LP*3-0-3 etc. according to the species. Since *LP*3 and *ASR* are members of the ABA/WDS induced protein superfamily *(Chang et al., 1996; González and Iusem, 2014; Padmanabhan et al., 1997)* care was taken to ensure that all sequences retrieved contained the ABA/WDS domain using PFAM v.31 *(Bateman and Finn, 2007; Mistry et al., 2007; Schaeffer et al., 2017)*. Sequence names and accession numbers used in phylogenetic tree reconstruction according to species is shown in the Supplementary Table 1. Supplementary Table 2 represents majorly represented species in which three or more ABA/WDS genes are present and its use if further detailed in section 2.3.

### 2.2 Phylogenetic analysis

The phylogenetic history of *LP*3 and *ASR* was determined using MEGA X (Kumar et al., 2018). Firstly, this was done by looking at the whole nucleotide sequences, then by looking only at the conserved ABA/WDS region of the sequences, the conserved N-terminal zinc binding region before the gymnosperm insertion, and then finally by looking only at the variable C-terminal NLS/DNA binding region of the sequences, producing a total of four different trees called FullSeq-tree, ABA/WDS-tree, N-tree and C-tree, respectively. The taxonomic phylogenetic tree was constructed using the Timetree software (Kumar et al., 2017).

For FullSeq-tree, sequences were aligned using the MUSCLE algorithm (Edgar, 2004), checked for errors and the multiple sequence alignment was exported for further analysis. The Maximum Likelihood (ML) Phylogenetic best fit model was determined in MEGA X by log-likelihood analysis of each model and the one with the highest AICc score was used. The resulting best fit model for the sequences used in tree FullSeq-tree was the Kimura 2 parameter model. The phylogenetic tree was created using the Maximum likelihood method combined with the Kimura 2 parameter model with a gamma parameter of 1, 30 with 1000 permutations.

ABA/WDS-tree was constructed by isolating the highly conserved ABA/WDS nucleotide domains from the MSA of *LP*3 and *ASR* and exporting those for phylogenetic analysis. The ML phylogenetic model was determined in the same manner as previously described, with the resulting best model by log-likelihood analysis being the Kimura 2 parameter model. The tree was constructed using the ML method with the Kimura 2 parameter with a Gamma parameter equal to 1,0846 with 1000 permutations.

Tree N-tree was constructed by extracting the N-terminal nucleotide region of *LP*3 and *ASR*. Due to the presence of incomplete sequences that did not cover this section of the gene, the following sequences were excluded from the phylogenetic analysis: **LP*3-0 Cupressus sempervirens; *LP*3-1 Pinus sylvestris; *LP*3-1 Pinus hwangshenensis; *LP*3-0 Pinus masssoniana; *LP*3-2 Pinus taeda; *LP*3-2 Pinus sylvestris* and all *LP*3-3 sequences. The ML phylogenetic model was determined in the same manner as previously described, with the resulting best model by log-likelihood analysis result being the Kimura 2 parameter model. The tree was therefore constructed using the ML method with the Kimura 2 parameter model with a gamma parameter equal to 1, 27 with 1000 permutations.

Tree C-tree was constructed by focusing on the C-terminal NLS/DNA binding regions of the sequences. The ML phylogenetic model was determined in the same manner as previously described, with the resulting best model by log-likelihood analysis result being the Kimura 2 parameter model. The following partial sequences were excluded from the phylogenetic analysis due to too short C-terminal sequences: **LP*3-0 Cupressus sempervirens, *LP*3-0 Pinus masssoniana; *LP*3-2 Pinus taeda; *LP*3-2 Pinus sylvestris* and all *LP*3-3 sequences. The tree was therefore constructed using the ML method with the Kimura 2 parameter model with a gamma parameter equal to 1, 5588 with 1000 permutations. Analysis of gene duplication events and construction of the corresponding gene duplication tree was also carried out in MEGA X, using FullSeq-tree as a template on which to perform the analysis.

### 2.3 GC% and RCSU analyses

Sequence names and accession numbers for determining GC and RCSU content according to species are shown in the Supplementary Table 2

The Relative Synonymous Codon Usage (RSCU) is a measure of the codon usage bias for a particular amino acid. As such codon bias is can be a measure of how efficient and accurate a given gene translation is. The RSCU of the ABA/WDS genes found in majorly represented species (species in which at least three different ABA/WDS genes are present) was extracted using MEGA X. The average RSCU values per amino acid were then calculated and used to determine which codon was on average the most used in a particular gene. Codons with RSCU values above 1 are abundant, whilst codons with RSCU values below 1 are less abundant. Codons for Methionine and Tryptophane were not included since these amino acids are encoded by only one codon. Stop codons were not included either as they are only involved in transcription termination and therefore not in transcription efficiency.

Individual gene parameters from majorly represented species such as total GC, GC1, GC2, GC3 contents, Codon Adaptive Index (CAI), and Effective Number of Nucleotides (ENC) were computed using CAICal (http://genomes.urv.cat/CAIcal/). The CAI is derived from the codon usage of highly expressed genes in organisms and is positively correlated with transcription levels. As such the CAI is often used as a proxy for expression levels (Sharp and Li, 1987). The ENC is a number corresponding to the overall codon bias in a given gene, with 20 symbolising a complete bias of only one codon per amino acid whereas a value of 61 symbolises a completely unbiased codon usage, with each available codon being used equally for a given amino acid (Wright, 1990). Statistical analyses were done in R using p=0,05 as the significance threshold. Gene parameters were first compared using a Levene test to ensure variance equality among the different genes, followed by either ANOVA or a Kruskal-Wallis test. If either of these indicated the presence of a significantlly different group then a Tukey HSD test was performed in the case of ANOVA and a pairwise Wilcoxon test in the case of a Kruskal-Wallis. Pearson correlations between the different gene parameters were also done. For this the different *ASR* and *LP*3 genes were grouped together.

### 2.4 Mode of evolution of different subsections of the ASR/LP3 gene

A codon-by-codon selective pressure analysis provides insights into which amino acids in a protein are undergoing selective constraints or not. This in turn allows one to suggest which amino acids are likely to change over time. Visualisation of codon selective pressure as defined by the ratio of synonymous to non-synonymous codons *ω = dN/dS* on the *LP*3-0 gene was done using the complete CDS of all *LP*3-0 homologous sequences were uploaded to the Selecton server for selective pressure analysis using the M8 model that allows for positive selection (Stern et al., 2007). The sequences were aligned in the server using the MUSCLE algorithm (Edgar, 2004) and the *Pinus taeda *LP*3-0* CDS was set as the reference query. Statistical analysis of the calculated selective pressure was also performed using Selecton with default settings by calculating the log likelihood ratio between M8 and the null model M8a. The same procedure was repeated for *LP*3-1, *LP*3-3, *ASR*1, *ASR*2, *ASR*3 and *ASR*4. *P.taeda *LP*3* and *S. lycopersicum ASR* sequences were set as references on which to visualise sites of selective pressure. There were not enough *LP*3-2 orthologous sequences for this analysis to be performed on it.

## 3. Results

### 3.1 Phylogenetic analysis of LP3 and ASR

On a broad level, it can be seen that *LP*3-1 is the most ancestral form of *LP*3, from which *LP*3-2 diverged, followed by *LP*3-0 and *LP*3-3. The *ASR* gene phylogeny show that *ASR*4 is the ancestral sequence in tomato, followed by *ASR*3, being *ASR*2 and *ASR*1 the result of are more recent diverge event (Figure 1). The clear distinction between the *ASR* and *LP*3 seems to correlate with the divergence of angiosperms and gymnosperms, around 313 MYA (Supplementary Figure 1, www.timetree.org).

**Figure 1:**
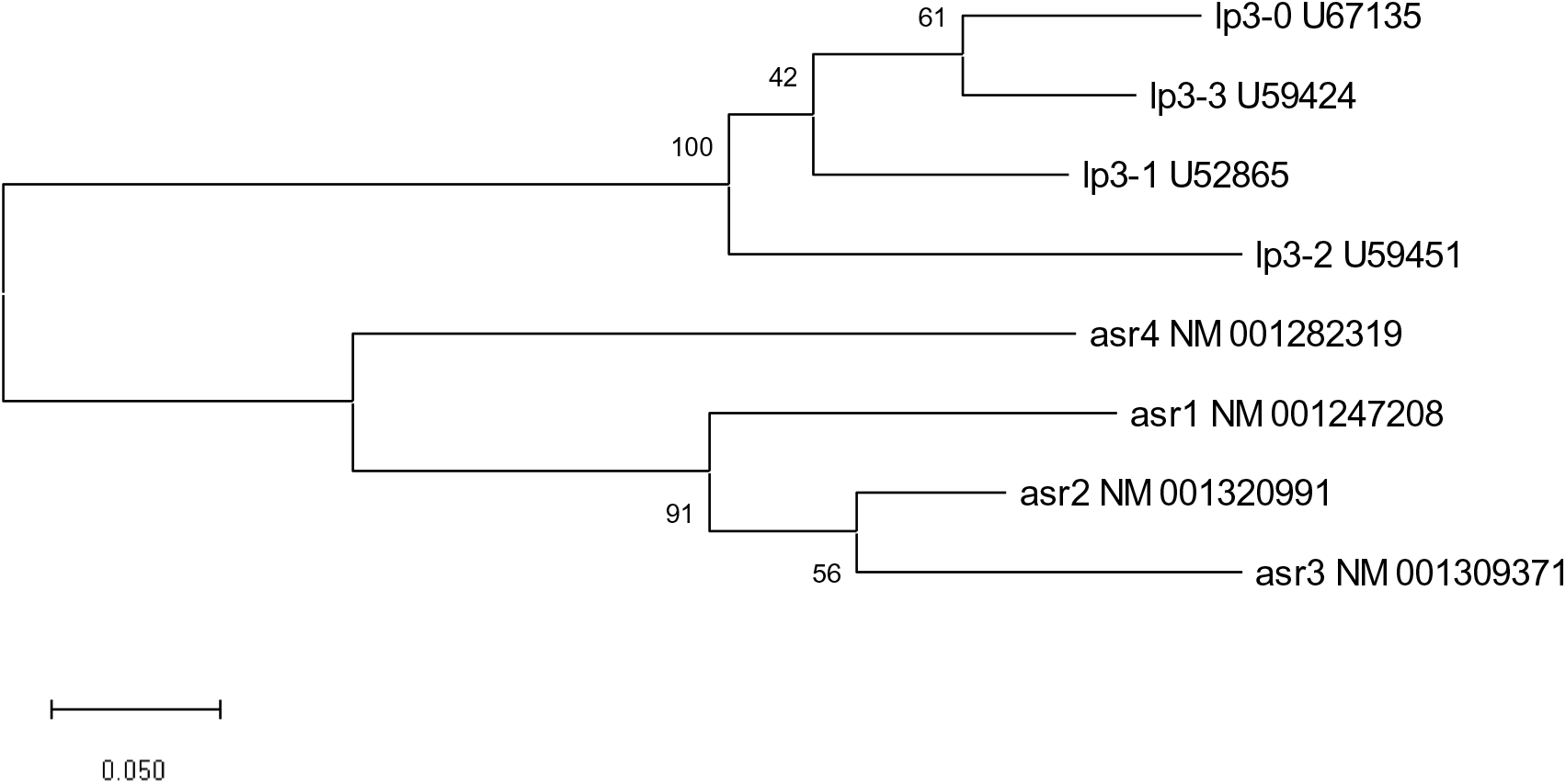
Phylogenetic tree of Pinus taeda LP3 and Solanum lycopersicum ASR sequences.

FullSeq-tree (*Figure 2*) shows that there is a clear divergence between the angiosperm *ASR* and the gymnosperm *LP*3 sequences. This could be to certain extent be attributed to the sequence insertion present between the N-terminal zinc binding domain and ABA/WDS regions in gymnosperm. Low bootstrap values within the gymnosperm nodes could be explained by the general high similitude between sequences which would result in them easily swapping positions during different bootstrap analyses. It can be observed that the *LP*3 sequences cluster together according to genus and isoform. There is also perfect clustering of *LP*3-1 and *LP*3-2 sequences within the *Pinus* cluster, however this is not the case when observing the *LP*3-3 cluster. In that instance, clustering of *LP*3-3 occurs with *LP*3-0-2 and *LP*3-0-3 of *Pinus sylvestris*. This might be indicative of orthology between *LP*3-3 and the *Pinus sylvestris* sequences shown. The grouping together of *Pinus taeda *LP*3-3* and *Pinus taeda *LP*3-0-2* with a very high bootstrap score could suggest that *LP*3-0-2 (PITA_00002958) might be actually a complete sequence of *LP*3-3. In *Picea* most of the sequences were annotated as *LP*3-0, except one sequence that was annotated as *LP*3-1 (MA45729). No *LP*3-2 and *LP*3-3 were available. The *ASR* sequences also show clear clustering together of *ASR*1, *ASR*2 and *ASR*3 sequences. The *ASR*1 of *Oryza sativa*, *ASR*3 of *Zea mays*, *ASR*2 of *Vitis vin*ifera and *ASR*3 of *Populus trichocarpa* show signs of divergence from the other *ASR* sequences. There are big genetic distances between the different *ASR* clusters, indicative of substantial divergence between the *ASR* genes in the angiosperms.

**Figure 2:**
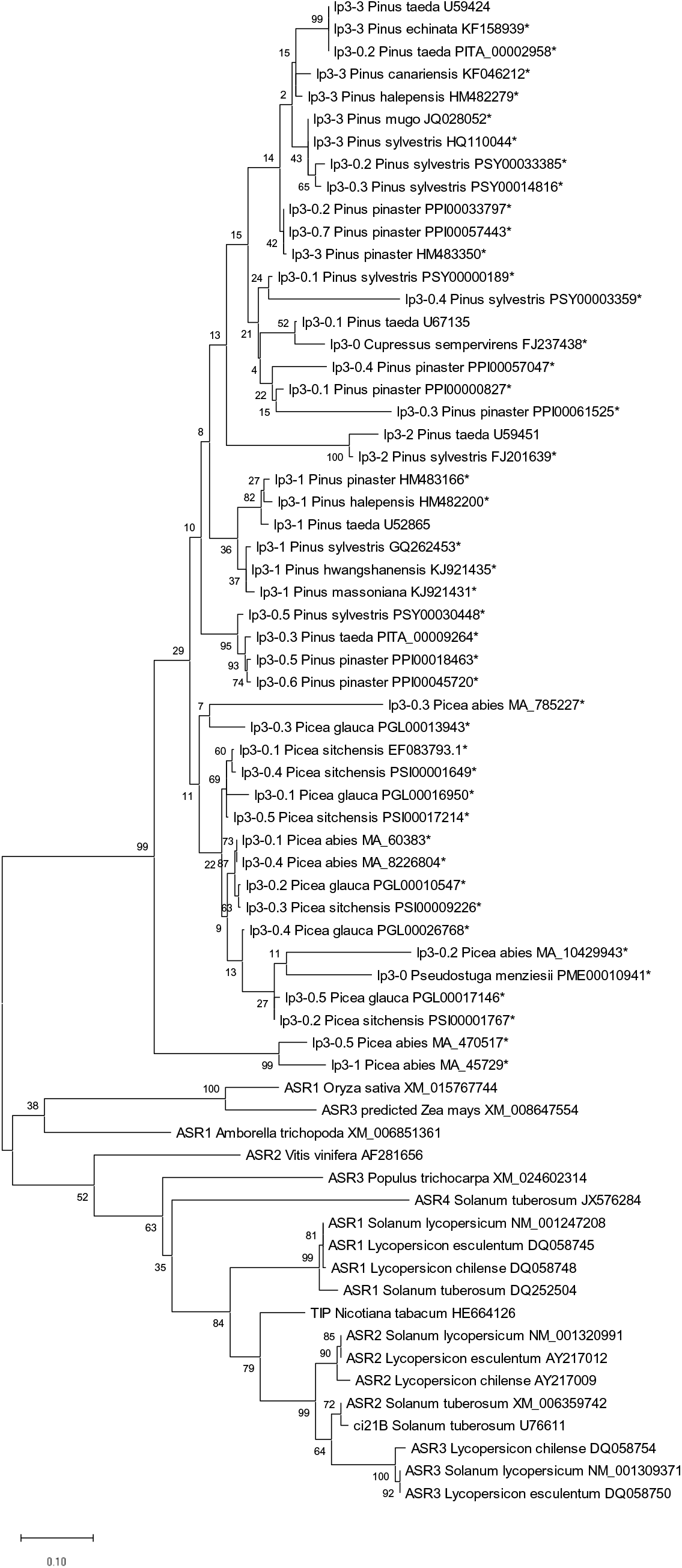
FullSeq-tree corresponding to the entirety of available CDS of ASR/LP3 sequences. Constructed in MEGAX using the ML method and the K2 model + gamma =1,2882. Genetic distance is given in number of substitutions per site. Bootstrap values are

Looking at ABA/WDS-tree (Figure 3) which focuses on the central ABA/WDS domain, there is yet again another clear grouping of angiosperm and gymnosperm sequences together. Within the gymnosperm grouping one can observe the sub-grouping of *LP*3-1 and *LP*3-2 together, while similarly to FullSeq-tree, the *LP*3-3 sequences are again broadly grouped together with some *LP*3-0 sequences grouped alongside them. The angiosperm sequences also show clear clustering of *ASR*1, *ASR*2 and *ASR*3 sequences with the *Oriza sativa ASR1* and *Zea mays ASR3* seeming to diverge from the other *ASR* sequences. As in FullSeq-tree, there is also a clear clustering according to genus, with *Picea* and *Pinus* sequences clustering together, respectively.

**Figure 3:**
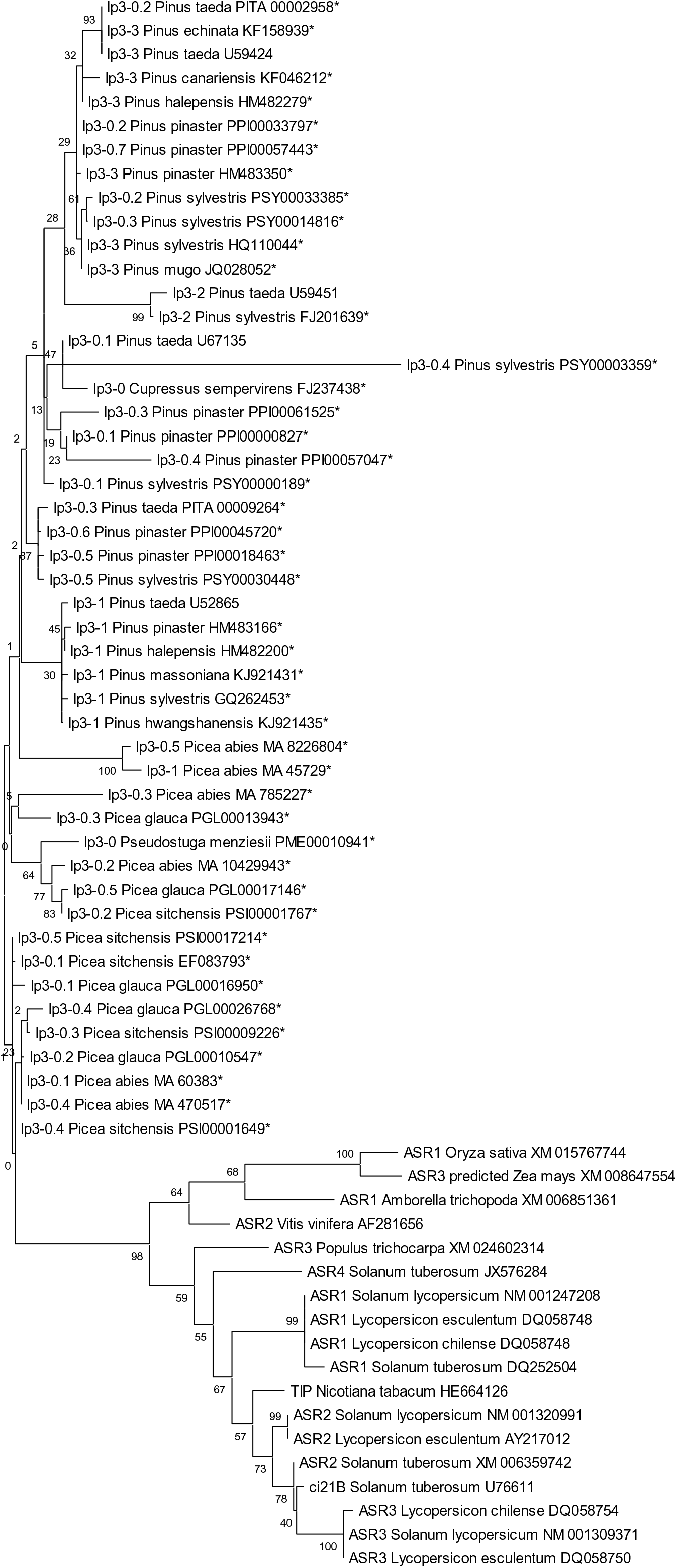
ABA/WDS-tree focusing on the conserved ABA/WDS domain within each gene. Constructed in MEGAX using the ML method and the K2 model with gamma=1,0846. Genetic distances are in number of substitutions per site. Bootstrap values are shown next to each node

N-tree focuses on the N-terminal region of *ASR* and *LP*3, to which zinc ions are theorised to bind (Figure 4), here it can be observed that there is not a definite separation between angiosperm and gymnosperm sequences, with *ASR*2 from *Vitis vinifera* and *ASR*4 from *Solanum tuberosum* being the most divergent sequences when focusing on the N-terminal region. *LP*3-1 sequences in *Pinus* formed a different cluster although with low support. After this node, *ASR* and *LP*3 sequences mostly segregate according to their order. Like the trees previously analysed, the *ASR* sequences cluster according to *ASR*1, *ASR*2 and *ASR*3, with the *Oriza sativa ASR1* and *Zea mays ASR3* diverging together. In contrast to previous trees, there is a less clear clustering of *LP*3 sequences according to genus. There remains however a very short genetic distance between each of the *LP*3 sequences, indicative yet again of a high level of conservation between the sequences.

**Figure 4:**
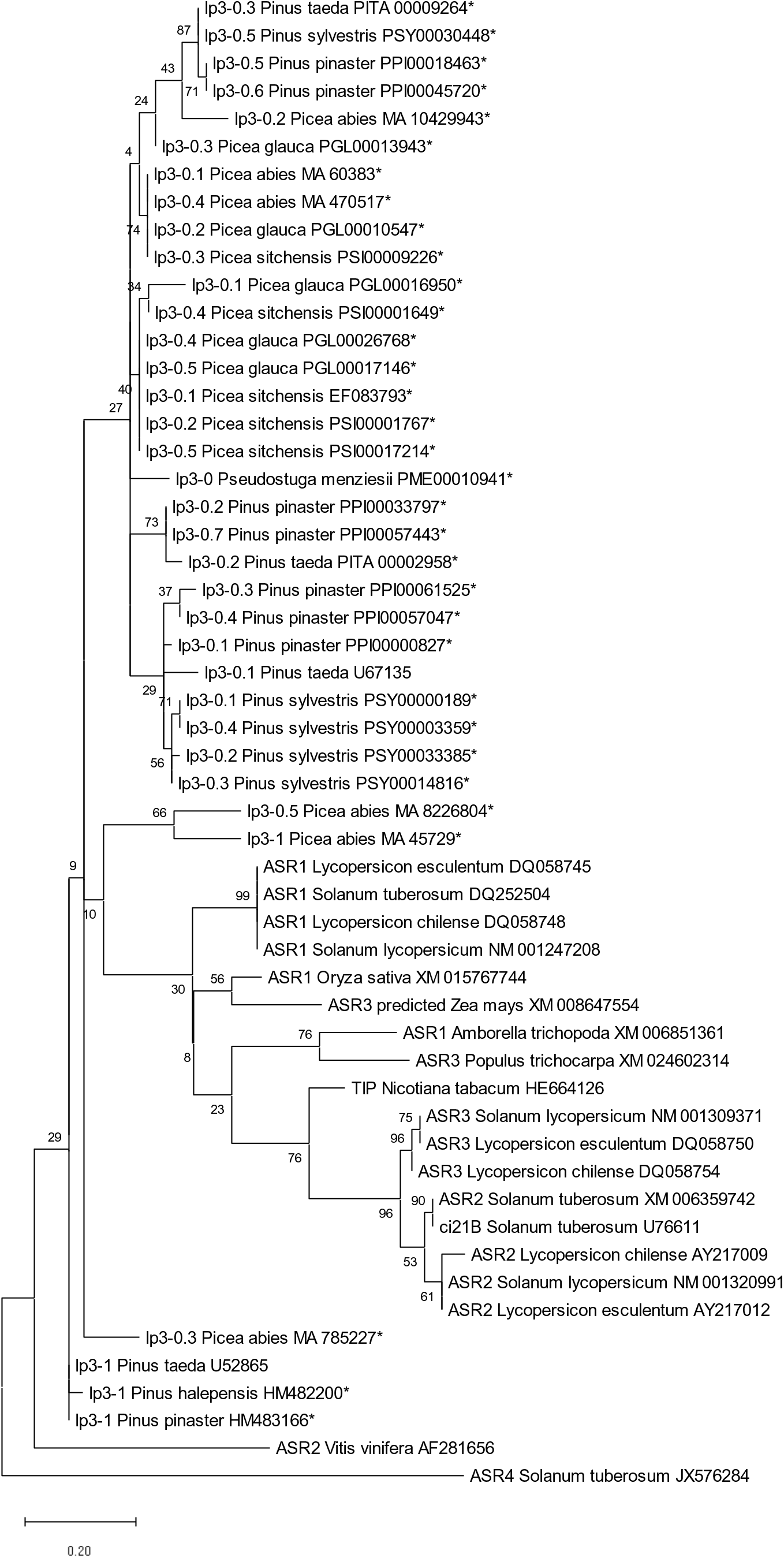
N-tree focusing on N-terminal of ASR and LP3 genes. Constructed with MEGAX and using the ML methodand K2 model with gamma =1,3701. Genetic distances are in number of substitutions per site. Bootstrap values are shown next to each node.

C-tree, which focuses on the variable C-terminal regions of *ASR* and *LP*3 which contains the NLS/DNA binding domain (Figure 5), again shows partial segregation of *ASR* and *LP*3 sequences. Here the tree is rooted by the *ASR*4 of *Solanum tuberosum*, **LP*3-0-3 Pinus pinaster* and *Picea abies *LP*3-0-2* and *LP*3-0-3 sequences, indicative that these sequences are the most divergent in the C-terminal NLS region. Subsequently, the *ASR* and *LP*3 sequences do segregate according to their order (similar to what can be observed for the other trees). Within the *ASR* cluster the *ASR*1, *ASR*2 and *ASR*3 groups show low genetic distances, which suggests that the C-terminal region of those sequences are well conserved. The *LP*3 C-terminal sequences also do not show much divergence between them as seen by the small genetic distances between the sequences. The exception are **LP*3-0-3, *LP*3-0-4 and *LP*3-0-2* sequences of *Picea abies*, which show a high level of divergence from the other *LP*3 sequences. The segregation within the *LP*3 sequences showed a similar clustering mostly according to the genus.

**Figure 5:**
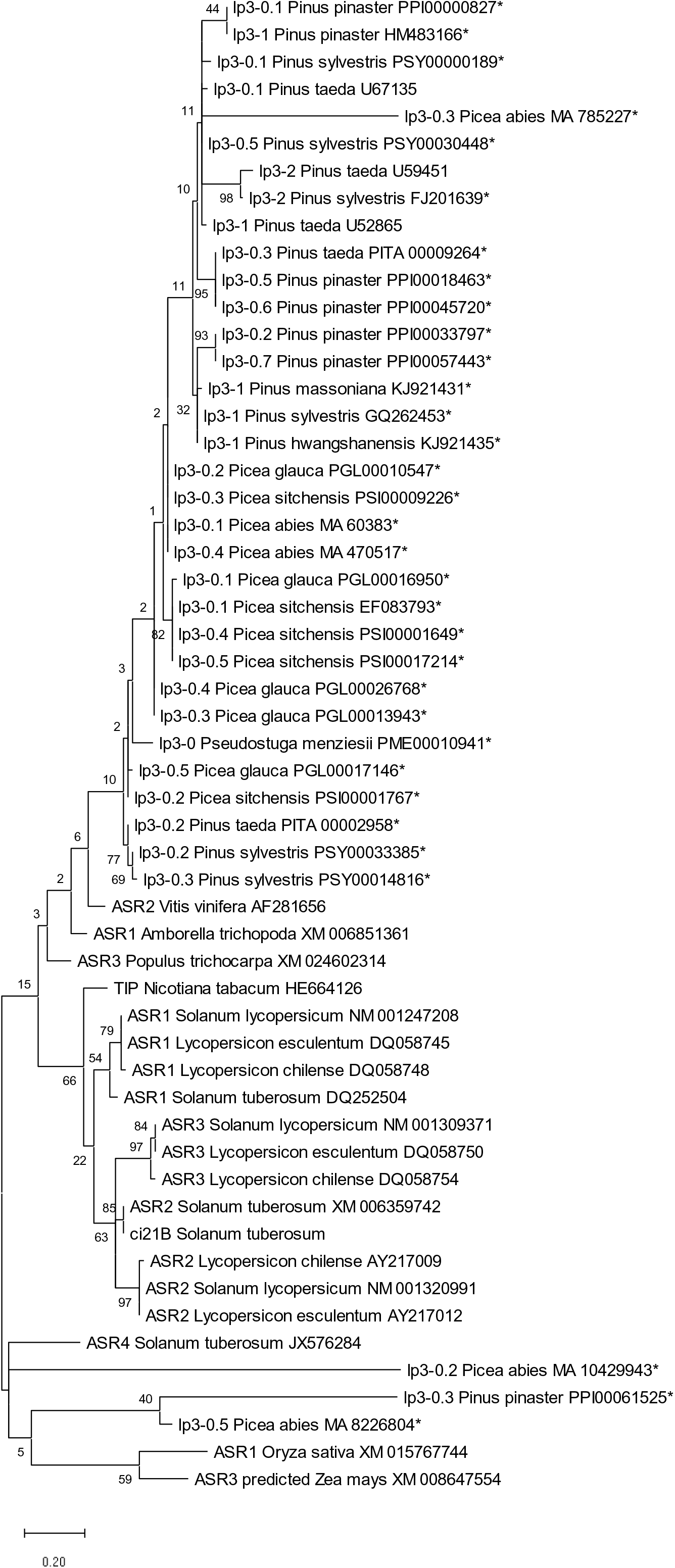
C-tree focusing on the C-terminal region of ASR and LP3 genes. Constructed with MEGA X and using the ML method with K2 model and gamma= 1,5588. Genetic distances are in number of substitutions per site. Bootstrap values are shown next to each node

### 3.2 Gene parameters and RSCU analyses

There appear to be differences in average preferential codon usage between the different ABA/WDS genes (Table 1, Figure 6). Both *LP*3_0 and *LP*3_3 have the same most highly used codon AGG, encoding arginine, at RSCU values 3,71 and 6 respectively. *LP*3-2 and *ASR*2 both preferentially use CCA, encoding proline, at RSCU values 4 and 2,77 respectively. Furthermore, *ASR*2 has a second codon AGC, encoding serine, at RSCU 2,77. *LP*3_1, *ASR*3 and *ASR*4 have all got codons encoding for serine as their most used codons with codons UGC, AGC and AGU at RSCU values 3,86, 3,27 and 2,8 respectively. There were similarities in least used codons, such as the arginine encoding CGG and leucine encoding CUA.

**Figure 6:**
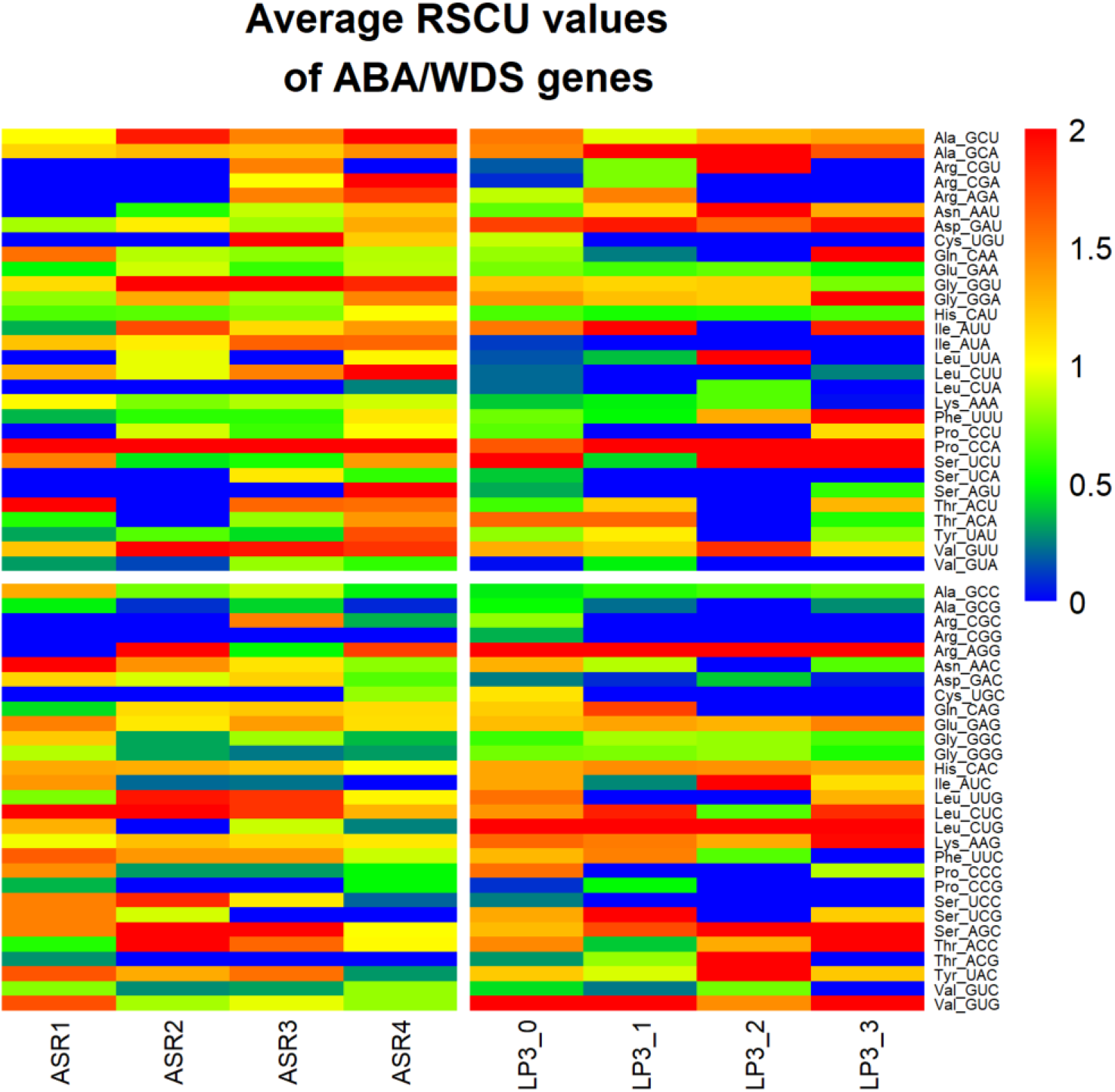
Heatmap of average codon usage per ABA/WDS genes. Codons separated according to whether they are AU or GC ended. Average codon usage above 1,5 are indicated in red while codon usage lower than 0,5 are indicated in blue. Produced in R.

**Table 1:**
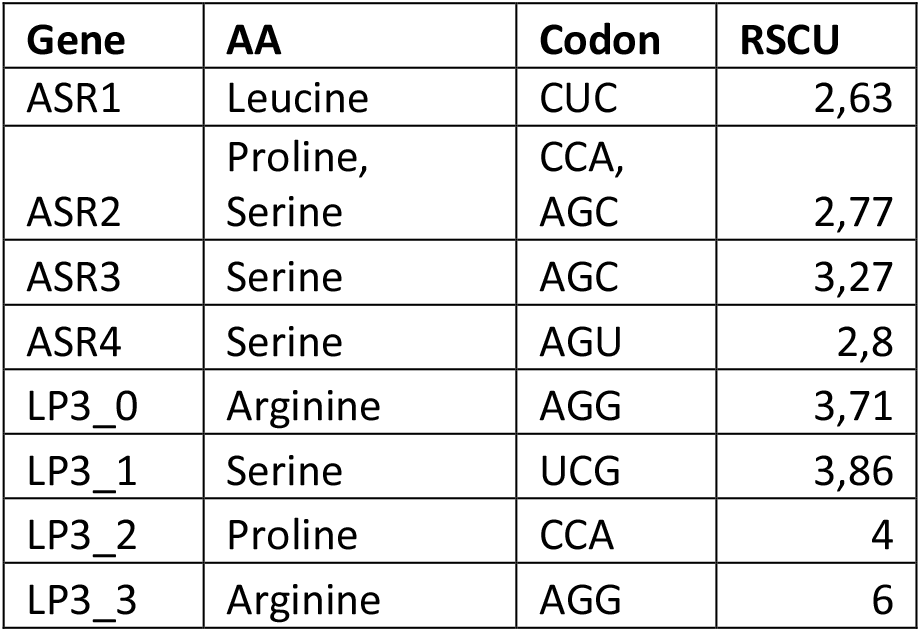
Average most used codon in ABA/WDS genes.

In all cases, the average GC2% of the ABA/WDS genes was lower than both average GC1% and GC3%. With the exception of *ASR*1, all ABA/WDS genes had on average higher GC1% than GC3% (Table 2). Levene test results showed that only GC1 and GC2 respected the equal variance among groups assumption for ANOVA (p>0,05). Total GC, GC3, ENC and CAI were thus tested using a Kruskal-Wallis test. GC1, GC2 and CAI had significant differences between the ABA/WDS genes (p<0,05) whereas total GC, GC3 and ENC showed no significant difference between the ABA/WDS genes (p>0,05).

**Table 2:** Average gene parameters of ABA/WDS genes.

| Gene | Length(bp) | CAI | GC%tot | GC1% | GC2% | GC3% | ENC |
| --- | --- | --- | --- | --- | --- | --- | --- |
| ASR1 | 323,25 | 0,8428 | 56,825 | 71,25 | 27,2 | 72,05 | 44,775 |
| ASR2 | 336 | 0,852 | 55,9 | 69,125 | 31,8 | 66,7 | 37,725 |
| ASR3 | 337,5 | 0,866 | 56,425 | 69,05 | 33,1 | 67,175 | 41,9 |
| ASR4 | 629 | 0,8217 | 43,46666667 | 56,5 | 36,6 | 37,33333333 | 50,03333333 |
| LP3_0 | 435,7 | 0,8472 | 50,69333333 | 64,76333333 | 30,43666667 | 56,89 | 51,02333333 |
| LP3_1 | 300,75 | 0,8528 | 51,25 | 68,975 | 27,875 | 56,975 | 42,4 |
| LP3_2 | 306 | 0,8435 | 51,6 | 71,95 | 29,35 | 53,5 | 53,9 |
| LP3_3 | 272 | 0,9073 | 52,83333333 | 67,73333333 | 35,43333333 | 55,43333333 | 37,33333333 |

Analysis of GC1 (Figure 7) suggests that on average *ASR*1 and *LP*3_2 have a higher GC1% than *LP*3_0, which itself has a higher GC1% than *ASR*4. *ASR*2, *ASR*3, *LP*3_1 and *LP*3_3 do not have significantly different GC1% from *ASR*1, *LP*3_2 and *LP*3_0.

**Figure 7:**
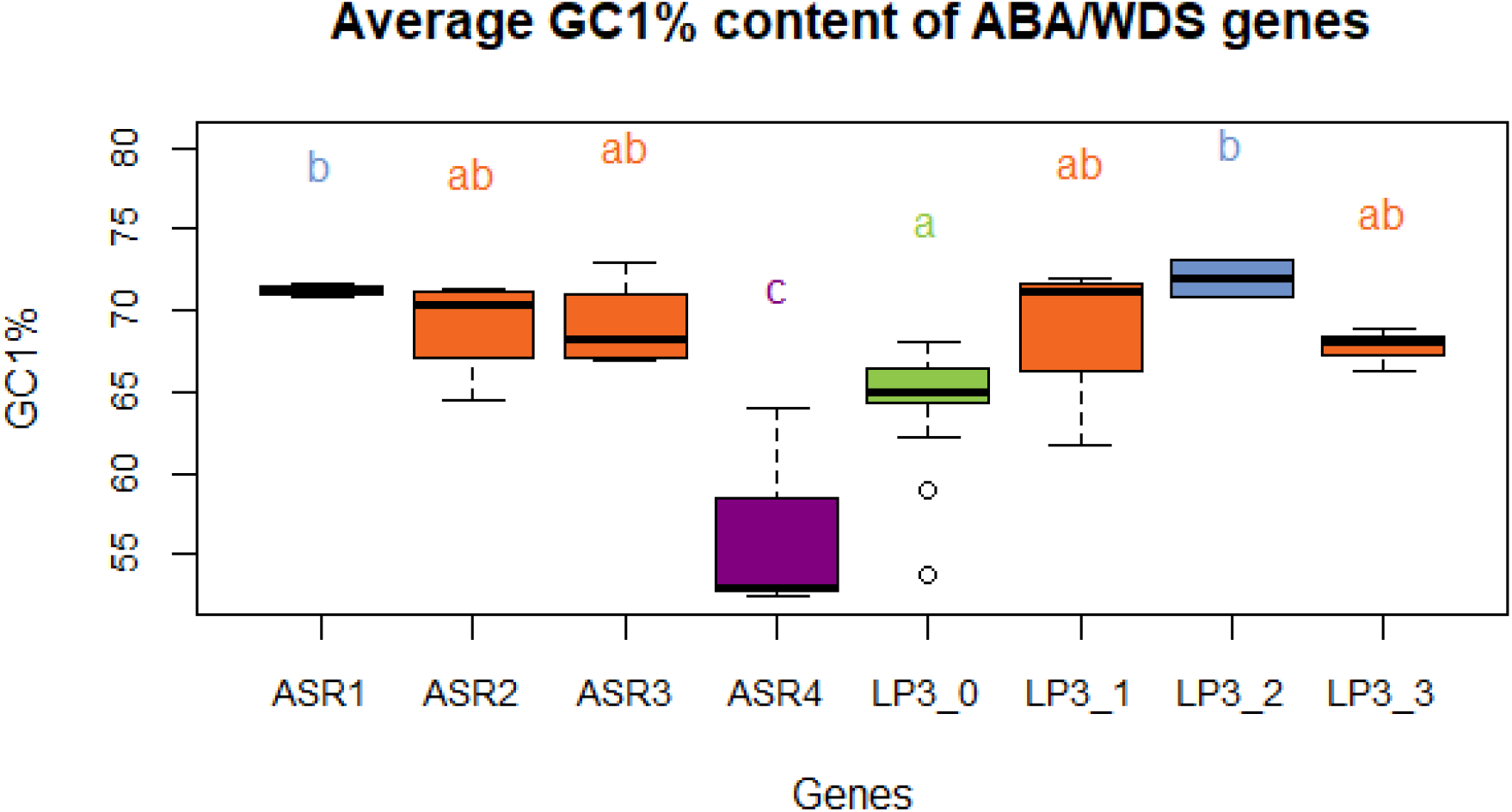
Comparison of the average GC1% per ABA/WDS gene, done in R, using a one-way ANOVA followed by Tukey HSD test.

Analysis of GC2% (Figure 8) suggests that *ASR*4 has on average a significantly higher GC2% than *ASR*1, *LP*3_0, *LP*3_1 and *LP*3_2. In turn, this analysis suggests that on average *LP*3_3 has a significantly higher GC2% than *LP*3_0. By contrast, *ASR*2, *ASR*3 and *LP*3_2 do not appear to have significantly different GC2% from each other.

**Figure 8:**
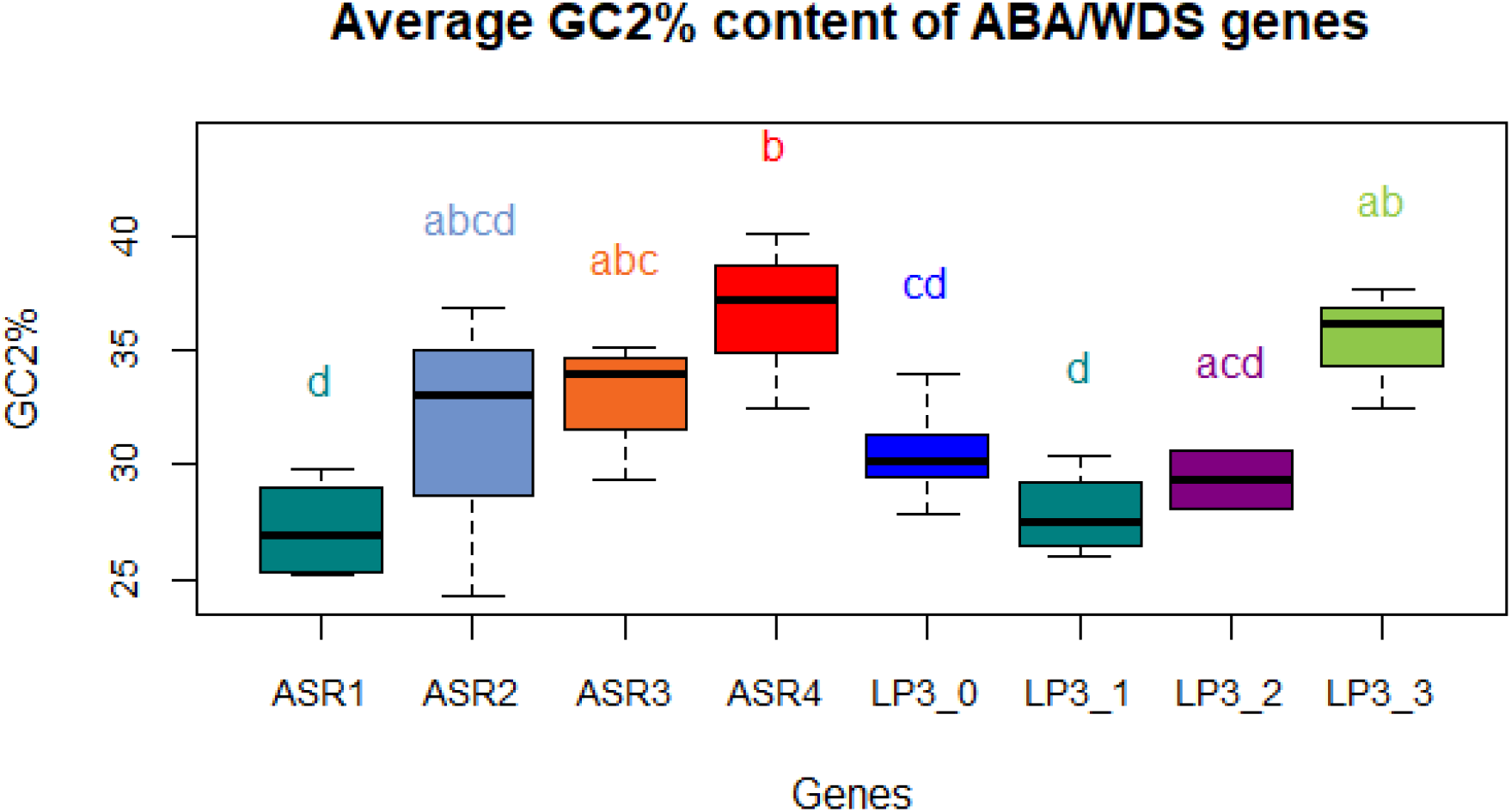
Comparison of the average GC2% per ABA/WDS gene, done in R, using a one-way ANOVA followed by Tukey HSD test.

There is also variation in the average CAI of ABA/WDS genes between the different majorly represented species (Figure 9). *Picea abies, Picea glauca, Populus trichocarpa, Solanum lycopersicum* and *Solanum tuberosum* have similar CAI levels to each other and are significantly lower than *Oryza sativa, Picea sitcchensis, Pinus pinaster, Pinus sylvestris, Pinus taeda* and *Zea mays*. *Oryza sativa* appears to have a significantly higher CAI than all other species apart from *Picea sitchensis, Pinus pinaster* and *Zea mays*. Finally, *Pinus sylvestris* appears to have a significantly lower average CAI than *Pinus pinaster, Zea mays* and *Oryza sativa*, but is significantly higher on average than *Picea abies, Picea glauca, Populus trichocarpa, Solanum lycopersicum* and *Solanum tuberosum* yet is not significantly different from *Picea sitchensis* and *Pinus taeda*.

**Figure 9:**
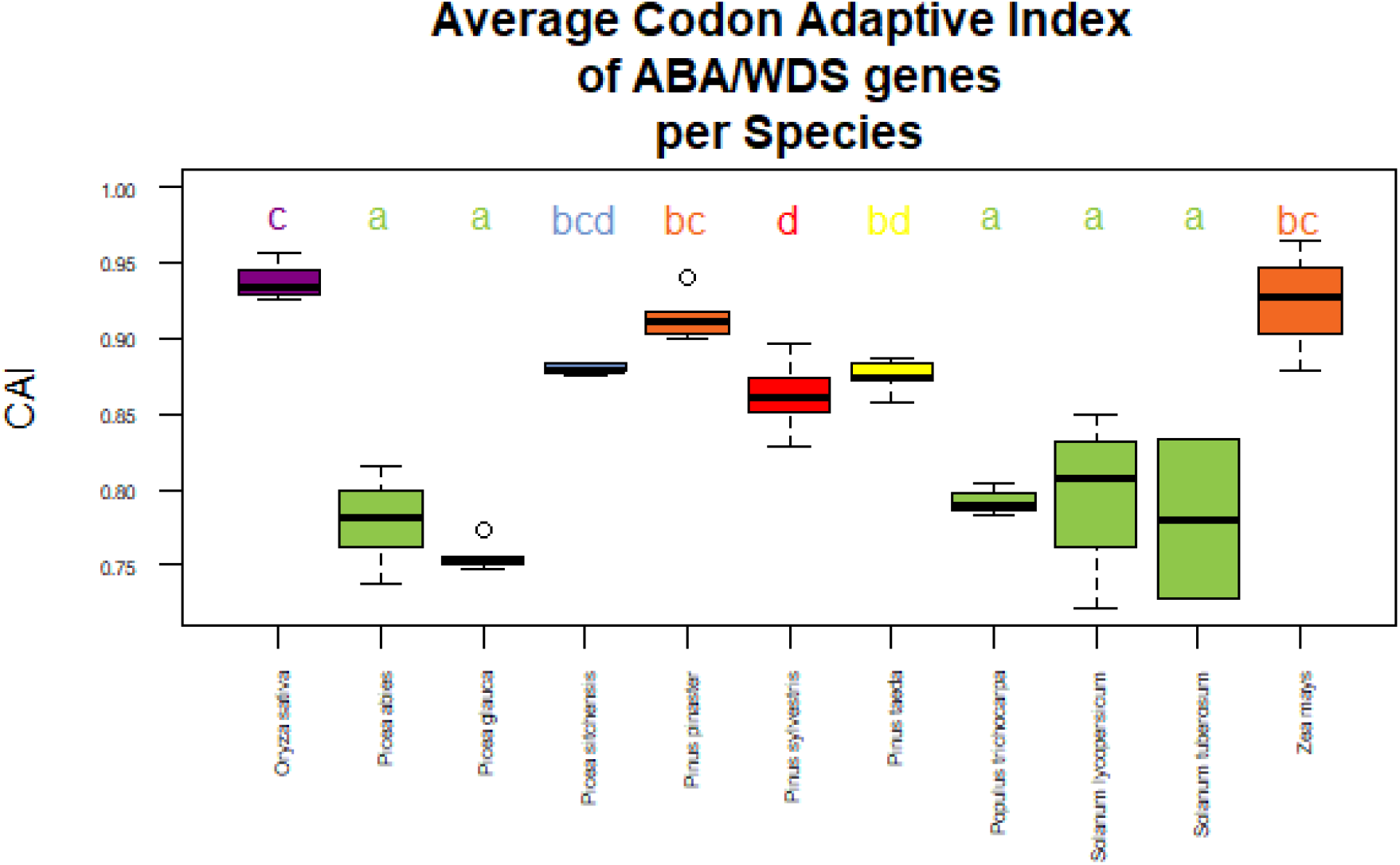
CAI per species of the ABA/WDS genes obtained from the values of majorly represented species, done in R, using a Kruskal-Wallis test followed by a pairwise Wilcoxon test.

Significant correlations were found between the different gene parameters in *ASR* and *LP*3 (Table 3; Table 4). In both *ASR* and *LP*3, gene length was negatively correlated with total GC%, GC1% and positively correlated with ENC. CAI was positively correlated with total GC% and GC3% in *ASR* and was positively correlated with GC1%. CAI was also negatively correlated with ENC in both cases. Total GC% was in turn positively correlated with GC1% and GC3% in both *ASR* and *LP*3 genes however total GC% was correlated with ENC in *ASR* only. GC1% was negatively correlated with GC2% in both *ASR* and *LP*3 and positively correlated with GC3% in *ASR* only. GC3% was negatively correlated with ENC in *ASR* but displayed no significant correlation with ENC in *LP*3.

**Table 3:**
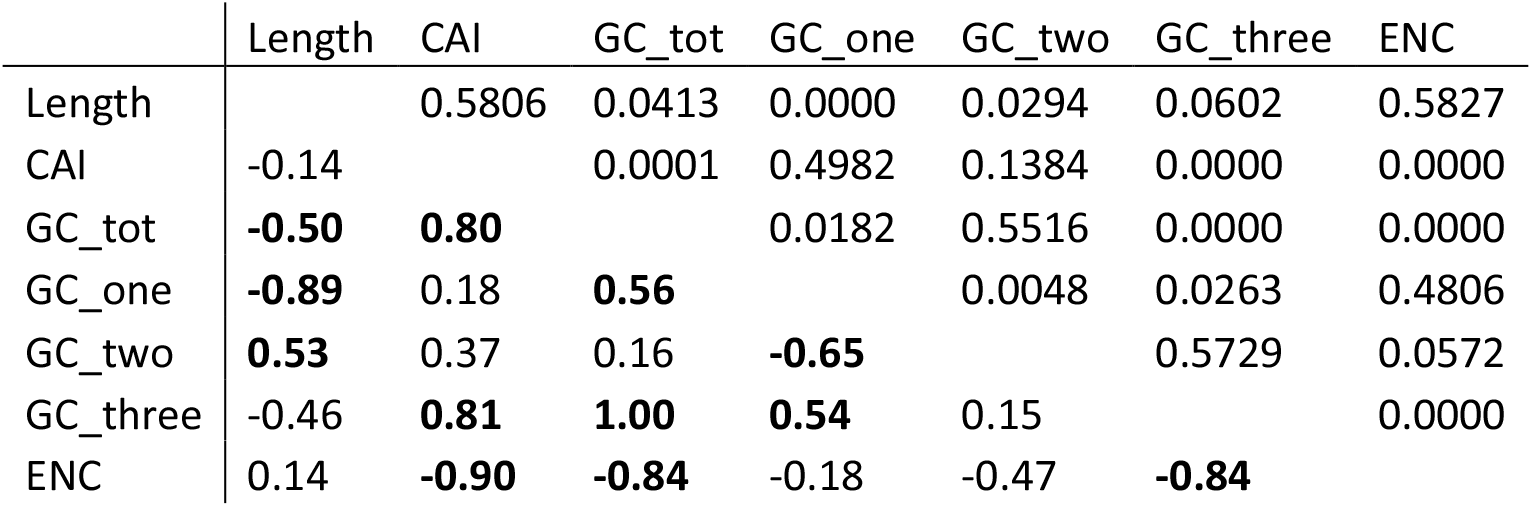
Pearson correlation matrix of ASR genes. Correlation values indicated below diagonal and p-values indicated above diagonal. Significant correlations shown in bold.

**Table 4.** Pearson correlation matrix of LP3 genes. Correlation values indicated below diagonal and p-values indicated above diagonal. Significant correlations shown in bold.

|  | Length | CAI | GC_tot | GC_one | GC_two | GC_three | ENC |
| --- | --- | --- | --- | --- | --- | --- | --- |
| Length |  | 0.5357 | 0.0284 | 0.0020 | 0.7432 | 0.6149 | 0.0018 |
| CAI | -0.10 |  | 0.1385 | 0.0431 | 0.7252 | 0.6568 | 0.0004 |
| GC_tot | **-0.35** | 0.24 |  | 0.0000 | 0.8198 | 0.0000 | 0.3053 |
| GC_one | **-0.48** | **0.33** | **0.58** |  | 0.0110 | 0.9112 | 0.1220 |
| GC_two | -0.05 | 0.06 | 0.04 | **-0.40** |  | 0.1903 | 0.5746 |
| GC_three | 0.08 | -0.07 | **0.65** | -0.02 | -0.21 |  | 0.5214 |
| ENC | **0.48** | **-0.54** | -0.17 | -0.25 | -0.09 | 0.11 |  |

### 3.3 Mode of evolution of different subsections of the ASR/LP3 gene

The reference sequences of *LP*3-0, *LP*3-1, *ASR*1, *ASR*2, *ASR*3 and *ASR*4 show signs of purifying selection, mainly focused on K, H and E residues, and no sites of positive selection (Figure 10, Figure 11, Figure 13, Figure 14, Figure 15, Figure 16; Supplementary Table 3,4,6,7,8,9). In contrast, *LP*3-3 presents signs of positive selection on residues N, S and T at positions 34, 37 and 61, respectively (Figure 12). These residues tend to turn into E, T and A, respectively (w-values = 2,8; 2,9; 2,9; p-values= 2, 1×10^−14^; 1, 2×10^−19^; 1, 4×10^−19^, Supplementary Table 5). On average, the ABA/WDS genes investigated here are all undergoing purifying selection, and all tend to similar amino acid sequences and codon usage as shown by the low Ka/Ks values between gene pairs (Table 5).

**Figure 10:**
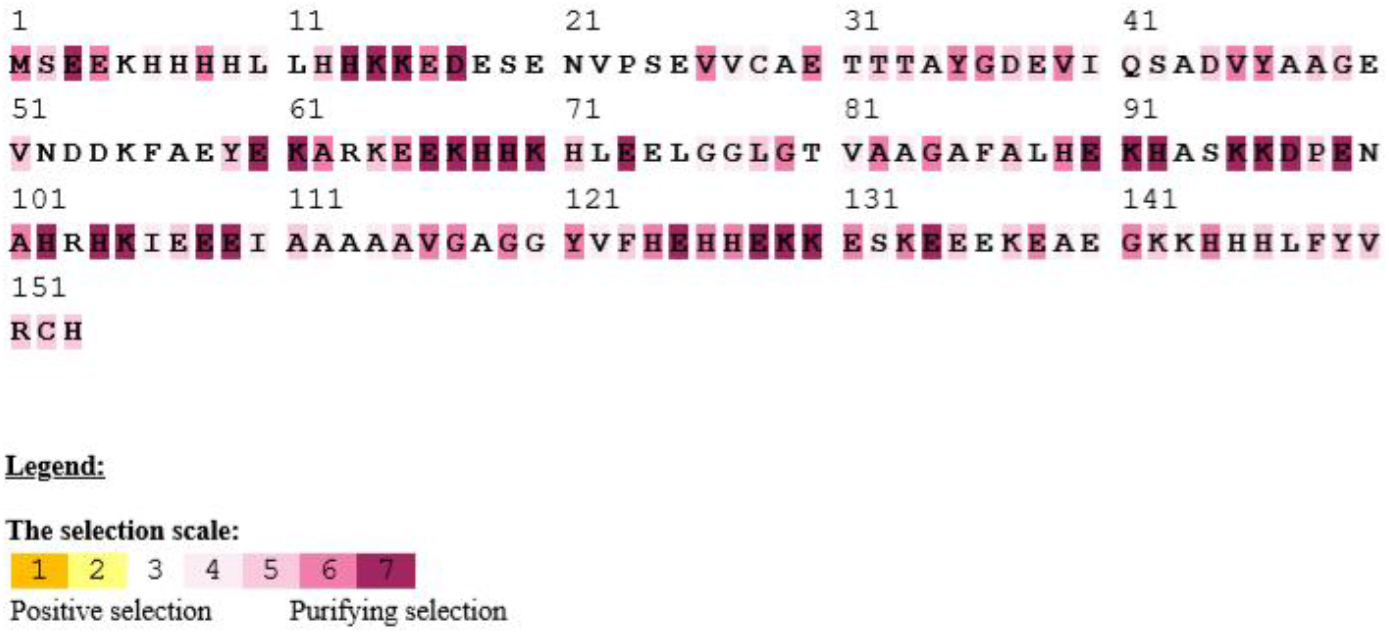
Selective pressures on LP3-0. Yellow colours indicate sites of positive/diversifying selection; purple colours indicate sites of purifying selection. Figure produced using Selecton.

**Figure 11:**
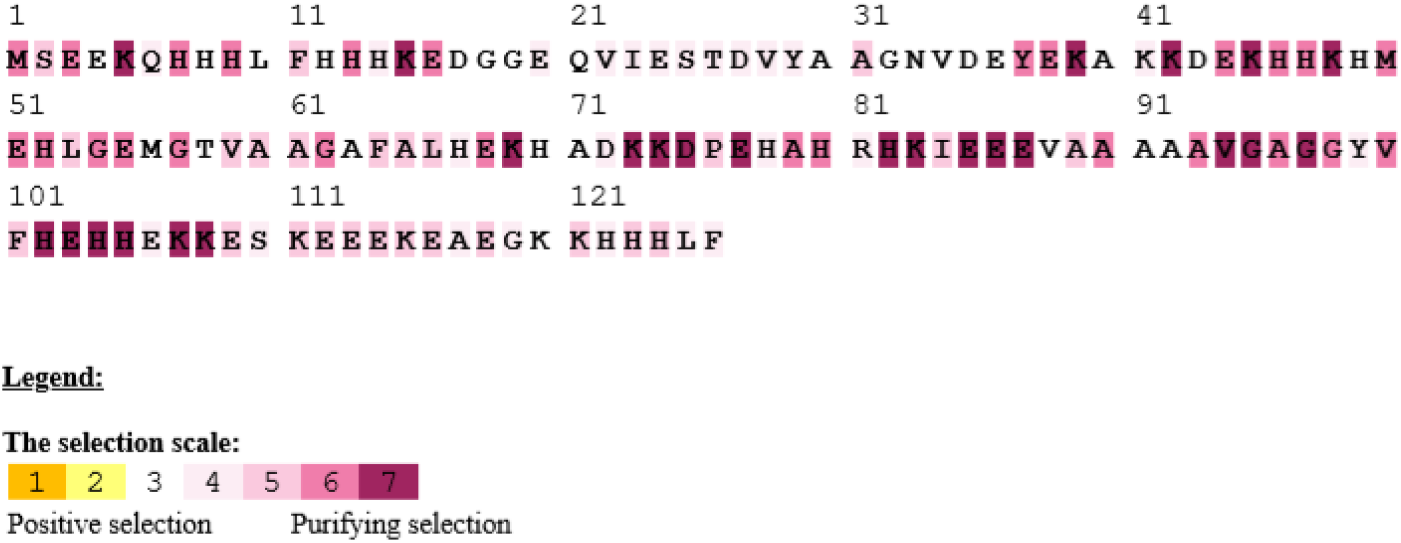
Selective pressures on LP3-1. Yellow colours indicate sites of positive/diversifying selection; purple colours indicate sites of purifying selection Figure produced using Selecton.

**Figure 12:**
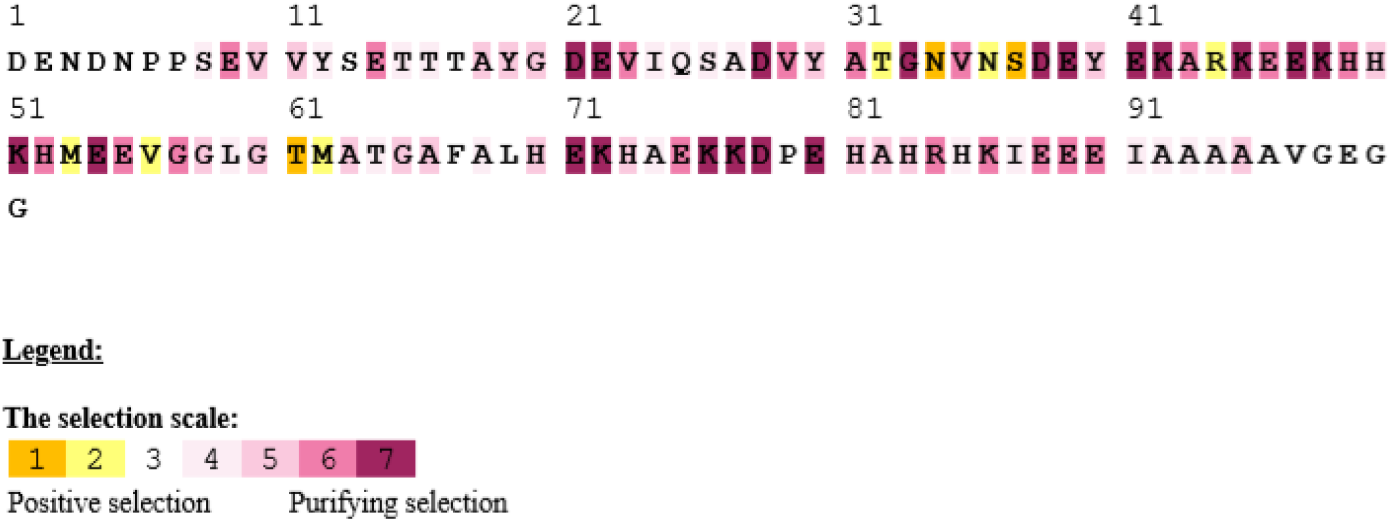
Selective pressures on LP3-3. Yellow colours indicate sites of positive/diversifying selection; purple colours indicate sites of purifying selection. Figure produced using Selecton.

**Figure 13:**
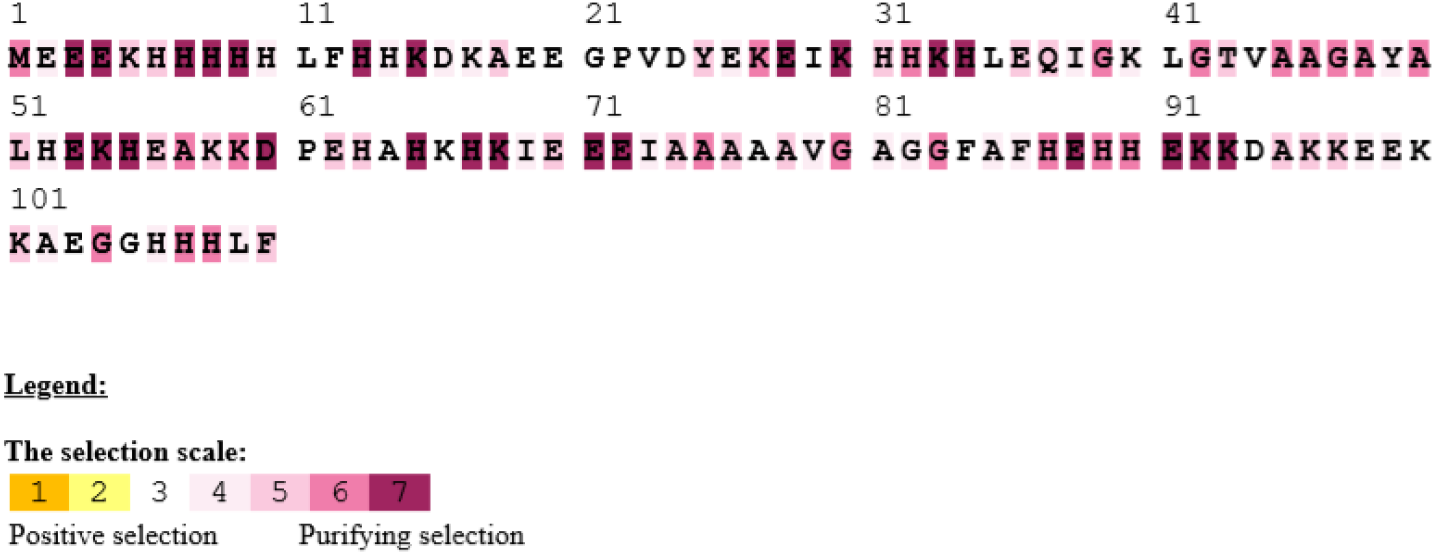
Selective pressures on ASR1. Yellow colours indicate sites of positive/diversifying selection; purple colours indicate sites of purifying selection. Figure produced using Selecton.

**Figure 14:**
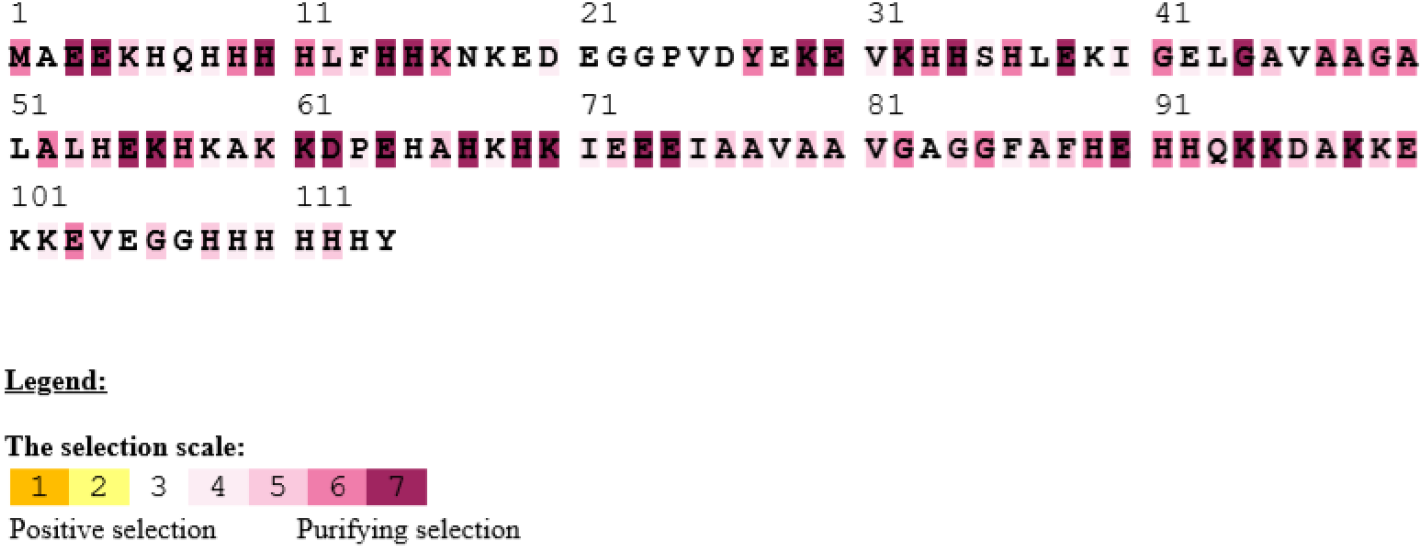
Selective pressures on ASR2. Yellow colours indicate sites of positive/diversifying selection; purple colours indicate sites of purifying selection. Figure produced using Selecton.

**Figure 15:**
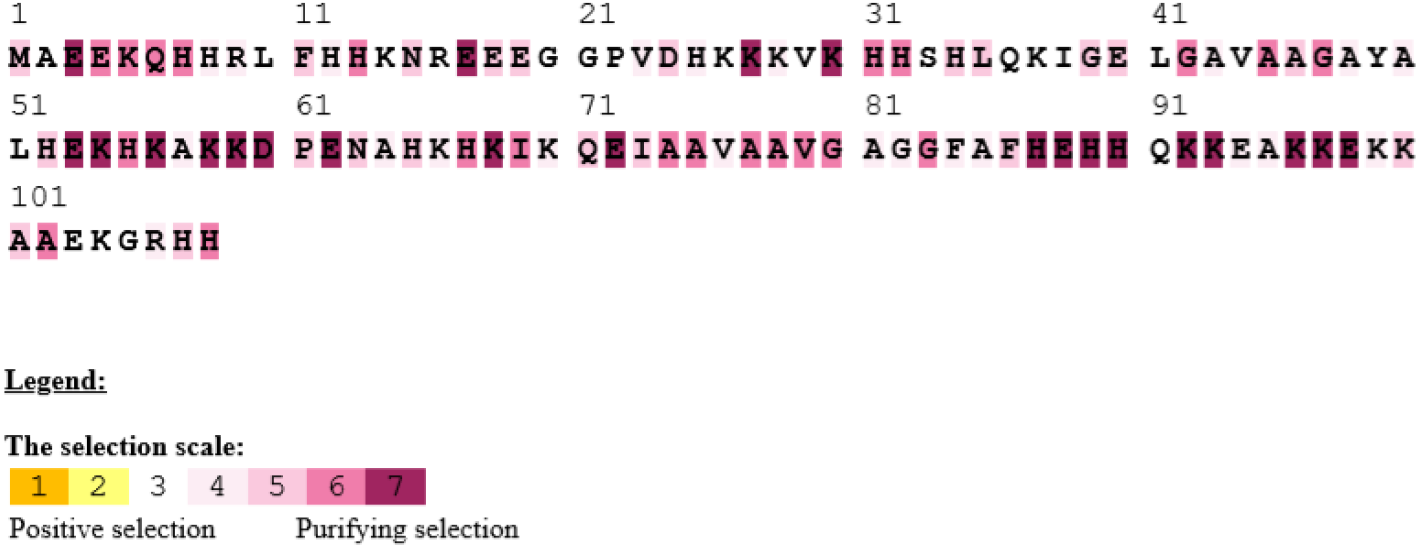
Selective pressures on ASR3. Yellow colours indicate sites of positive/diversifying selection; purple colours indicate sites of purifying selection. Figure produced using Selecton.

**Figure 16:**
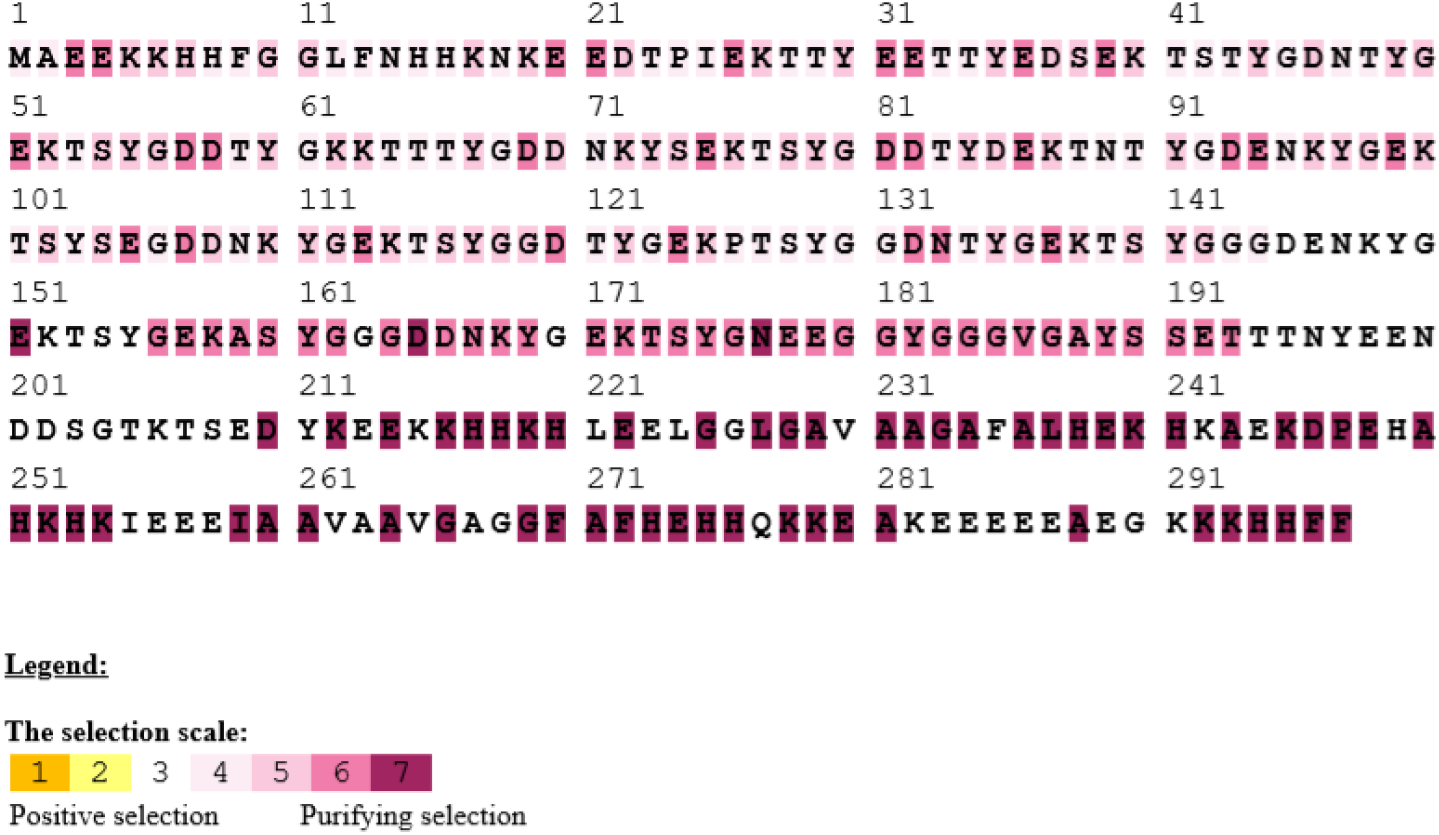
Selective pressures on ASR4. Yellow colours indicate sites of positive/diversifying selection; purple colours indicate sites of purifying selection. Figure produced using Selecton.

**Table 5:** 1 Ka/Ks estimates between (a) LP3 genes in Pinus taeda, and (b) ASR genes in tomato.

|  |  | LP3-0 | LP3-1 | LP3-2 | LP3-3 |
| --- | --- | --- | --- | --- | --- |
| U67135 | LP3-0 |  |  |  |  |
| U52865 | LP3-1 | 0.21 |  |  |  |
| U59451 | LP3-2 | 0.74 | 0.53 |  |  |
| U59424 | LP3-3 | 0.35 | 0.28 | 0.74 |  |

|  |  | ASR1 | ASR2 | ASR3 | ASR4 |
| --- | --- | --- | --- | --- | --- |
| NM001247208 | ASR1 |  |  |  |  |
| NM001320991 | ASR2 | 0.15 |  |  |  |
| NM001309371 | ASR3 | 0.25 | 0.06 |  |  |
| NM001282319 | ASR4 | 0.16 | 0.10 | 0.21 |  |

## 4. Discussion

### 4.1 LP3 genes are less divergent than ASR genes

While there is ample evidence that *ASR* and *LP*3 belong to the same gene family (González and Iusem, 2014), it is quite interesting that the nucleotide phylogeny reveals a clear segregation between the gymnosperm and angiosperm sequences, even within a highly conserved domain like the ABA/WDS domain. The separation between *LP*3 and *ASR* sequences are consistent with previous phylogenetic knowledge that angiosperms and gymnosperms diverged around 313 MYA (Barbara-Montoya et al 2018), between the Pennsylvanian and Permian periods, thereby giving the *ASR* and *LP*3 genes within both orders ample time to diverge (Hedges et al., 2015; Kumar et al., 2017).

The FullSeq-tree is corroborated by a previous study on *ASR* genes (Frankel et al., 2006). The phylogenetic analyses of different sub-sections of the *ASR*/*LP*3 genes provide insight into the different divergence rates occurring within each of them. The ABA/WDS-tree, which focuses on the ABA/WDS subsection, shows that there is very little divergence between the *LP*3 sequences, unlike the *ASR* sequences. This is indicative of a high level of similarity of the ABA/WDS domain within *LP*3 that is not seen in *ASR*. This can be explained by the action of negative selection acting on the *LP*3 sequences, or that the rate of substitutions in gymnosperms is lower than in angiosperms (Palmé et al 2009; Buschiazzo et al 2012; De La Torre et al., 2017).

This low level of phylogenetic divergence within *LP*3 can also generally be observed in the other two domain-based trees (N-terminal tree and C-terminal tree). The N-terminal tree focused on the putative zinc binding domains in the N-terminal region of *ASR*/*LP*3 genes, therefore it would make sense that there is a low level of divergence occurring since *ASR*/*LP*3 requires Zinc ions to adopt their functional conformation (Dominguez and Carrari, 2015; González and Iusem, 2014). A lack of clear segregation between *ASR* and *LP*3 can also be indicative of high purifying selective pressures thereby favouring a conserved nucleotide sequence. The lack of a definite separation between angiosperm and gymnosperm sequences is another argument in support of the essential and possibly similar role of the N-terminal domain in both clades.

C-tree, which focused on the putative NLS/DNA binding region of *ASR*/*LP*3, reveals similar topological properties as the N-tree but with a lower support to the branches discerning the different *ASR* gene family members. Both *ASR* and *LP*3 encode relatively small proteins and in the case *ASR*1 it has been previously shown that it does not require its putative NLS to enter the nucleus (Ricardi et al., 2012; Rom et al., 2006). This low level of divergence might indicate that the selective pressures focusing on these parts of the genes stem mostly from their role as transcription factors. Following this reasoning, it could be suggested that the sequence corresponding to *LP*3-0-3 *Picea abies* targets a different gene somewhere along the genome other than the *LP*3 gene family.

In all cases, it was observed that the *LP*3 genes had much lower divergence between them than the *ASR* genes. This is in line with previous research that found that coniferous genes have a low divergence rate among themselves when compared with angiosperm (Buschiazzo et al., 2012). In addition to all the individual reasons listed previously, previous transcriptomic analyses of homologous genes have shown that angiosperm genes tend to diverge much more strongly than coniferous genes (Li et al., 2010). Our phylogenetic analyses are therefore in line with previous known research.

### 4.2 Differences in GC1% and GC2% between genes, differences in CAI between species

While no significant difference was observed in total GC or GC3 content between the ABA/WDS genes, there were differences in the GC1 and GC2 contents between genes. In all cases the GC2 was lower than both GC1 and GC3, a pattern which has been observed in other gene analyses (Song et al., 2017, 2018). In all the major species investigated, CAI was on average highest in *Oryza sativa*, which in turn can be used to predict that ABA/WDS expression levels are highest within *Oryza sativa* (Sharp and Li, 1987). Previous research on *ASR* expression in rice has shown that it responds strongly to drought stress (Pérez-Díaz et al., 2014), so this high CAI is further indicative of high expression, which in turn lends credence to the other average CAI of ABA/WDS of the other investigated species being indicative of gene expression. It was interesting to see variations in CAI levels between the coniferous species, especially between the *Picea* and *Pinus* genuses, with *Pinus* species exhibiting higher CAI than *Picea* (except for *Picea sitchensis*). This in turn may be indicative that the ABA/WDS expression is higher in *Pinus* than in *Picea*, although a more thorough gene expression analysis within more species of each genus is required to confirm this hypothesis.

Previous gene studies have focused on the correlations of CAI with various other factors such as gene length, GC content and ENC (Gun et al., 2018; Song et al., 2017, 2018; Zhou and Li, 2009). Our study however showed no significant correlation between sequence length and CAI. CAI is often correlated with GC3% yet this correlation was only significant when concerning *ASR* genes and not the *LP*3 genes. Instead CAI was positively correlated with GC1% in *LP*3, which contrasts with previous gene studies (Gun et al., 2018; Song et al., 2017, 2018). Our study does reflect significant correlations between CAI and ENC within the ABA/WDS genes, indicative of a high translation efficiency found in many highly expressed genes (Sharp and Li, 1987; Wright, 1990)

### 4.3 ASR has a different codon usage than LP3

Another factor that contributes to differentiate *ASR* and *LP*3 gene families is the RSCU. Differences in codon usage were observed for several amino acids. Yet it was interesting to see that the amino acid with the highest codon bias was serine, although the individual codon with the highest bias was different among different genes. It has already been established that monocots and dicots differ in their codon usage for the homologous genes (Campbell and Gowri, 1990), therefore it is not improbable that gymnosperms would have different codon usages than angiosperms for homologous genes. Highly expressed genes have been shown to have both a more pronounced codon bias and higher overall GC content compared to lowly expressed genes (De La Torre et al., 2015; Song et al., 2017; Kuzniar et al., 2008), however no significant differences in overall average GC content between the ABA/WDS genes were observed in this study.

After gene duplication the original and new copies can go down multiple evolutionary paths (Innan, Kondrachov, 2011). While *LP*3-3 and *LP*3-0 genes have seemingly diverged recently they are affected by different modes of gene-duplication evolution. *LP*3-3 presents signs of positive selection on three residues and the rest are evolving either under neutral or purifying selection, whereas *LP*3-0 is only affected by purifying selection.

In *LP*3-3 those three residues (mutation) may have introduced new beneficial functional aspects to the original *LP*3 copy resulting in their fixation and maintenance through positive selection. This mode of gene-duplication evolution would suggest the acquisition of a novel function for *LP*3-3 gene. Notably purifying selection is acting on Lysine (K), Histidine (H) and Glutamate (E). *LP*3 and *ASR*, both belonging to the ABA/WDS gene family, are both highly hydrophilic protein groups therefore the presence of charged, polar amino acids is important in attaining their functional conformations(González and Iusem, 2014; Rom et al., 2006).

It has also been suggested that Zinc ions bind to lysine residues in the N-terminal region of both sets of genes, this binding being required for the proteins to finally obtain their functional conformation, thereby further explaining why mutations affecting Lysine are purified. Evidence of purifying selection can also be observed in both the ABA/WDS domain and C-terminal containing a putative NLS/DNA binding amino acid sequence, implying that both are important for the overall functioning of the genes. This is logical when considering that expressions of both *LP*3 and *ASR* are upregulated in presence of ABA and that both act as transcription factors in response to water-deficit stress (Padmanabhan et al., 1997; Wang et al., 2002).

### 4.4 Conclusion and future perspectives

Overall, this study highly suggests that while *ASR* and *LP*3 may have originated from the same common ancestor, they have undergone significant shifts in codon usage, maybe due to different evolutionary constraints. Different *ASR* genes have already been studied in depth, with *ASR*1 being the most substantially studied of them. This is not the case for *LP*3 for which more research opportunities are available. Further studies into *LP*3 mutants could produce similar phenotypes as observed in *ASR* mutants. Precise functions and genomic targets of *LP*3 could be hypothesized by homology with genes targeted by *ASR*. Precise mapping of the *LP*3 genes onto the *Pinus taeda* genome should also be done. Further research could establish the presence of paralogous genes of *LP*3 within *Pinus taeda* and their precise role in drought response. Since the responses of plants to cold stress are similar to their responses to drought stress, research into how *ASR* and *LP*3 are affected in terms of expression and cellular function in conditions of cold stress would also be an interesting research objective. Finally, a precise expression network between ASR/*LP*3 and downstream targets should be established since these have not yet been determined.

## Acknowledgments

We would like to thank Nicolas Delhomme and Sonali Ranade for bioinformatic support. We would like to acknowledge the UPSC Vinnova Center for Forest Biotechnology.

---

**Supplementary Table 1:**
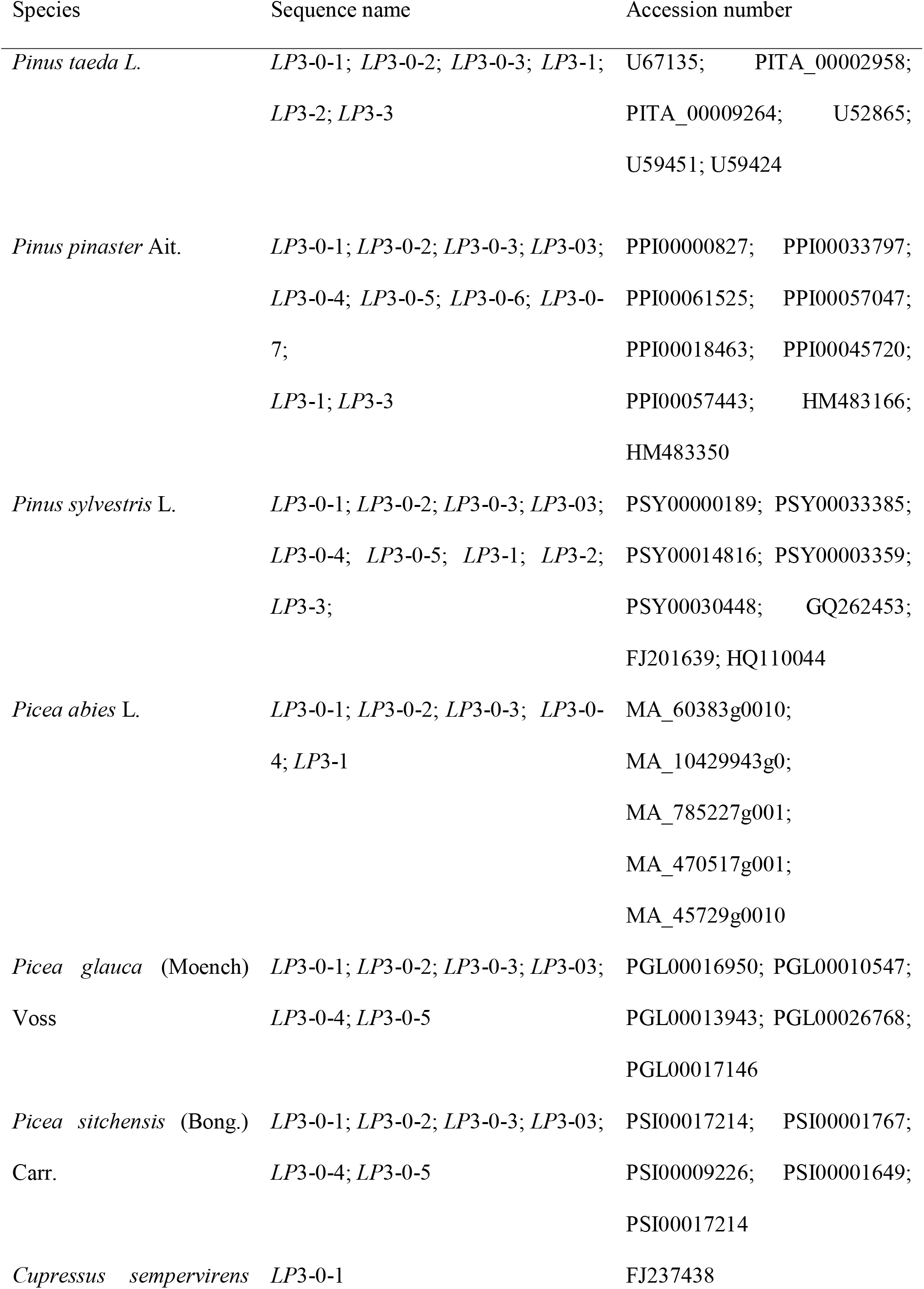
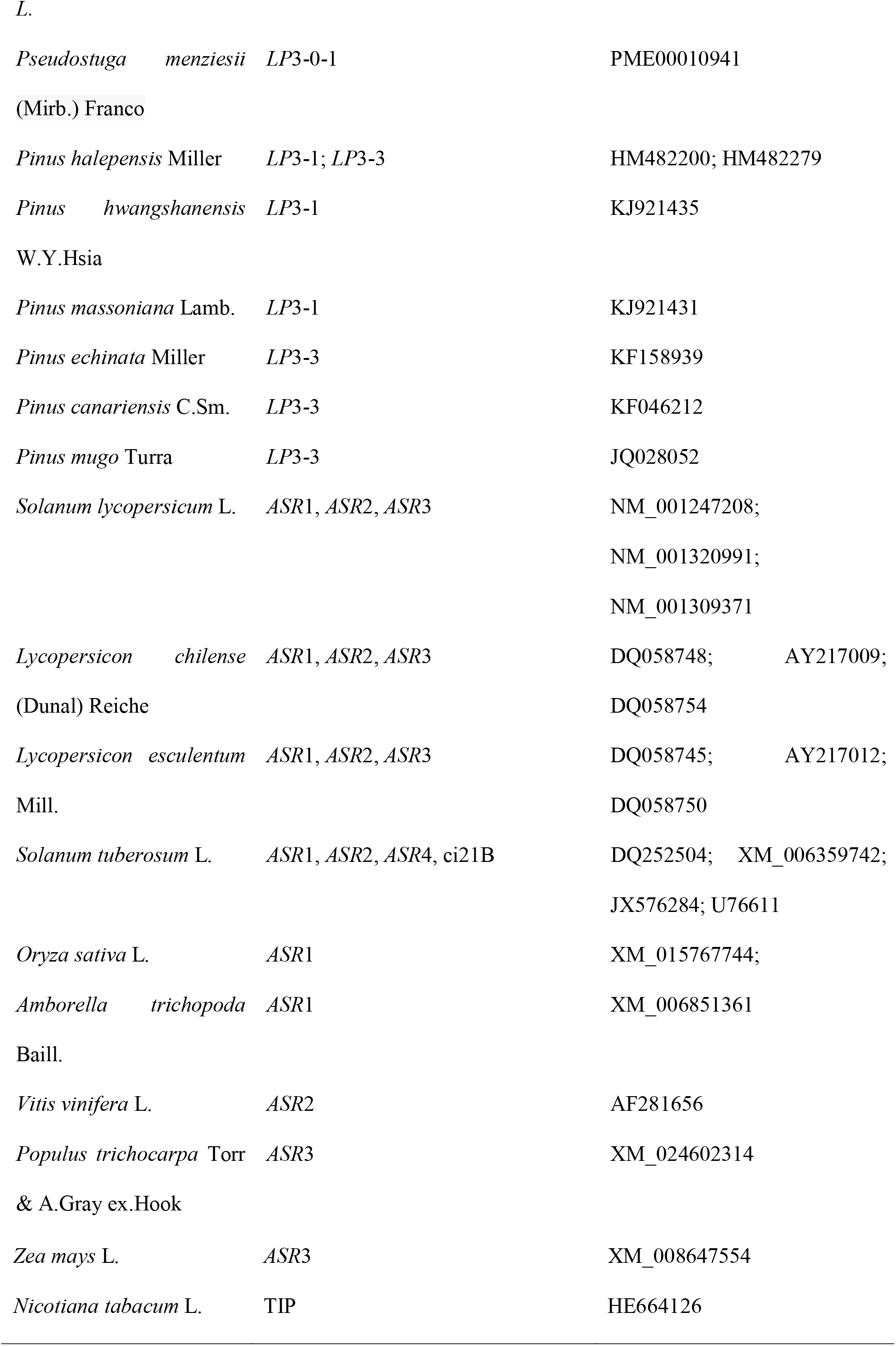
Sequence names and accession numbers used in phylogenetic tree reconstruction according to species.

**Supplementary Table 2:**
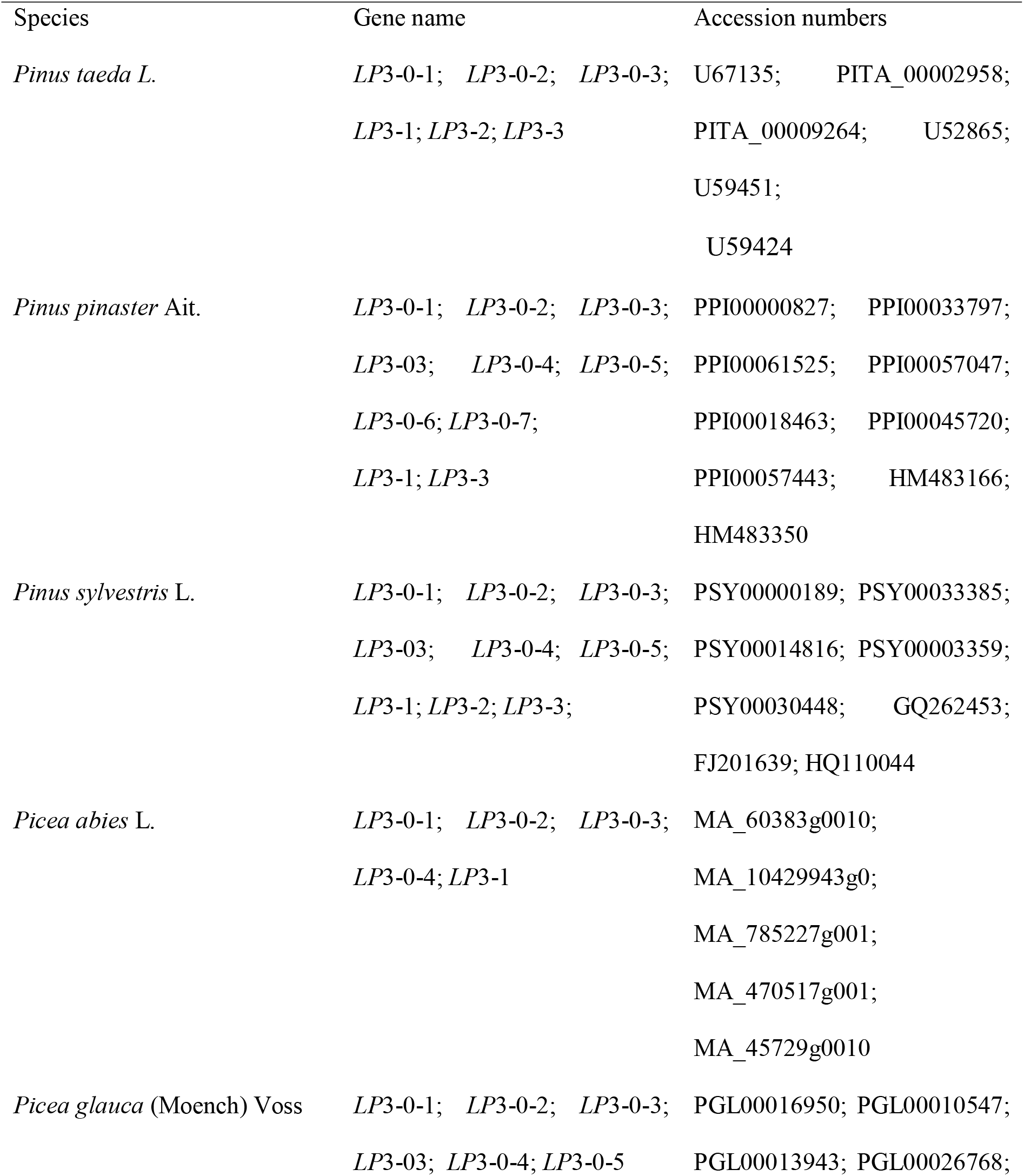
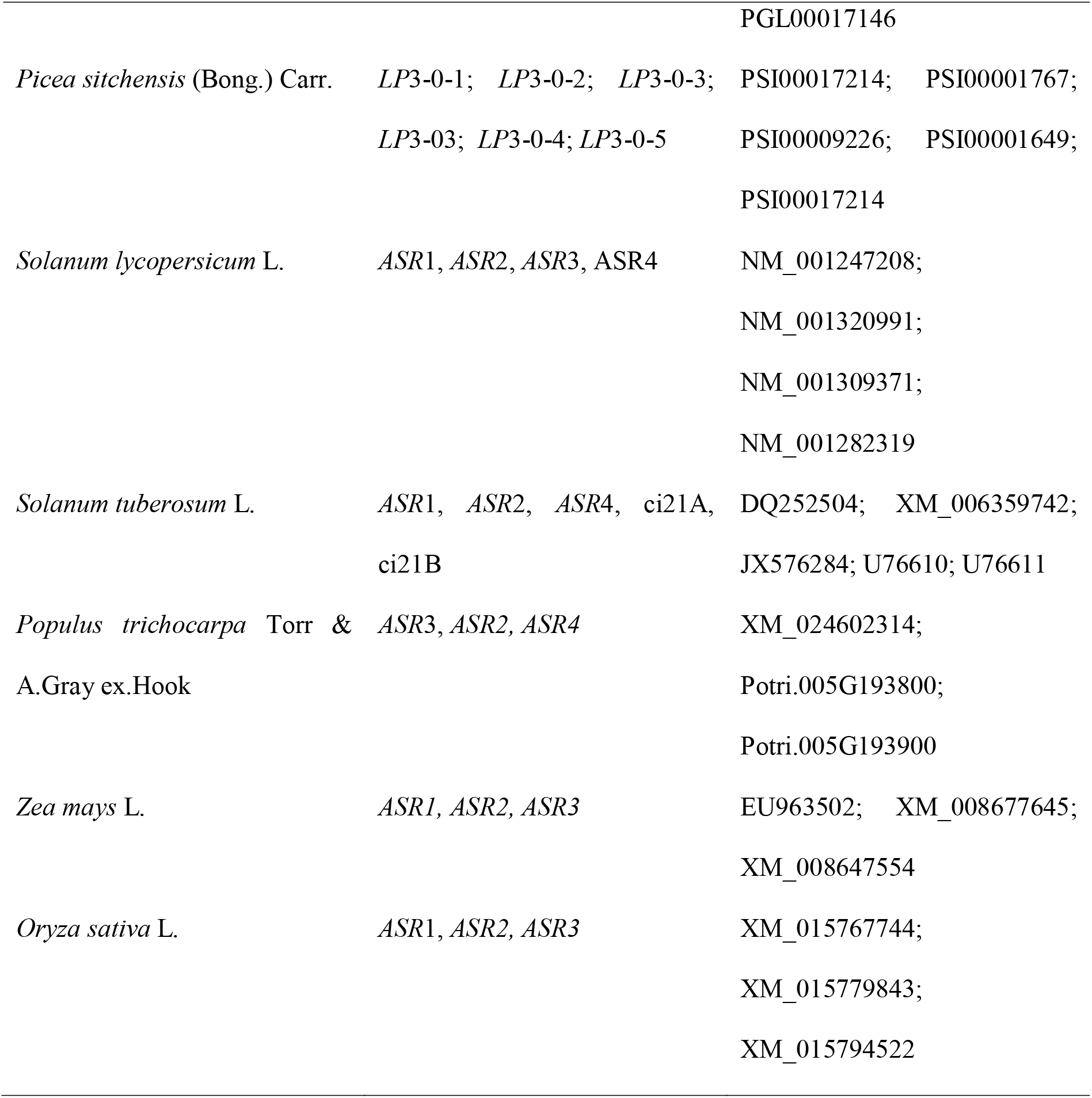
Sequence names and accession numbers from majorly represented species (i.e. species in which three or more ABA/WDS were found) for determining CAI, GC content, ENC and RSCU according to species.

**Supplementary Table 3:**
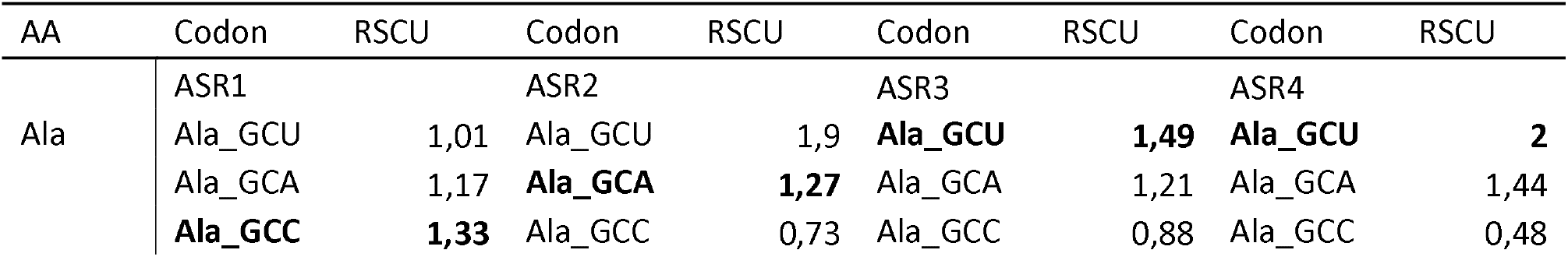
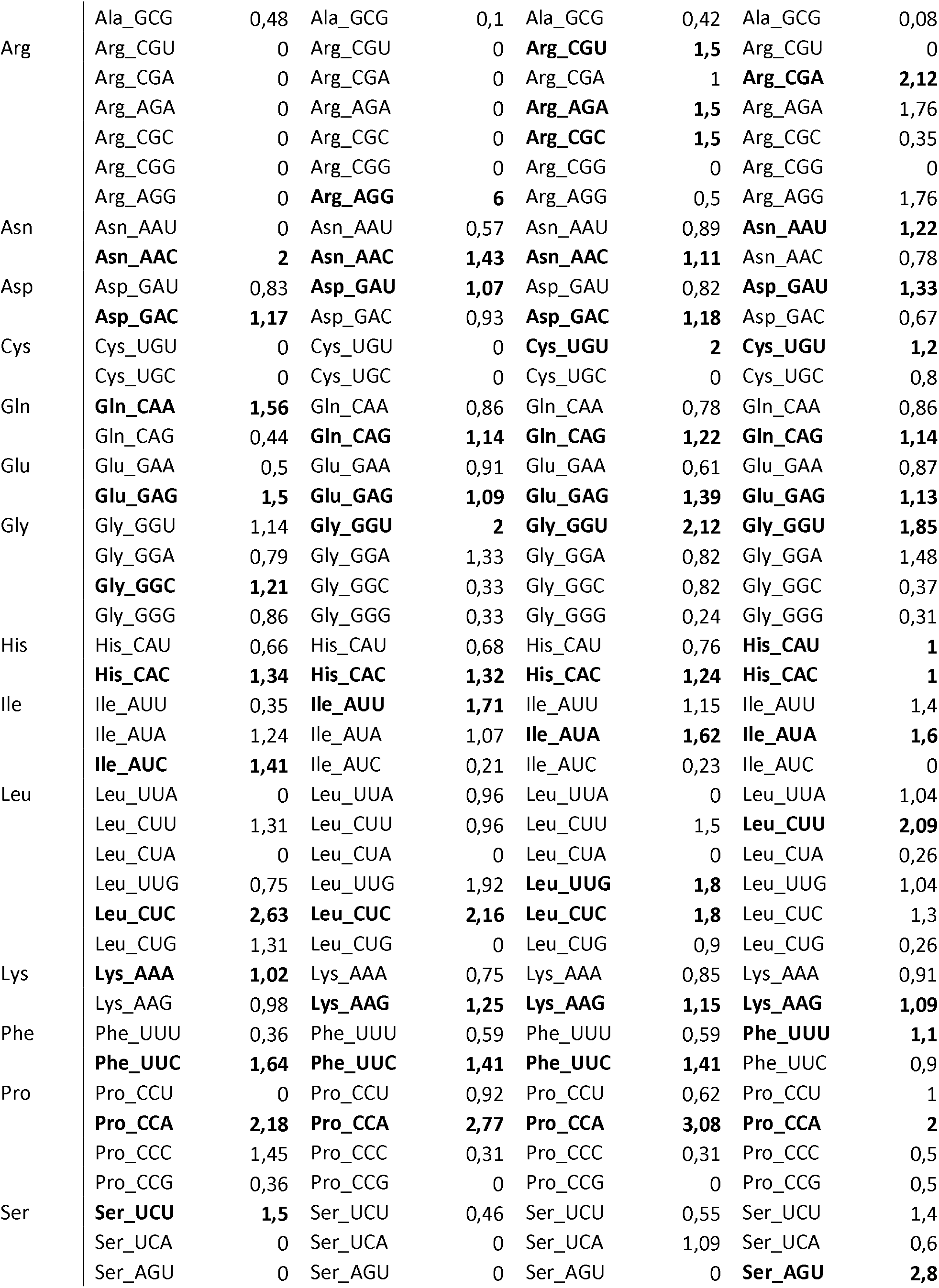
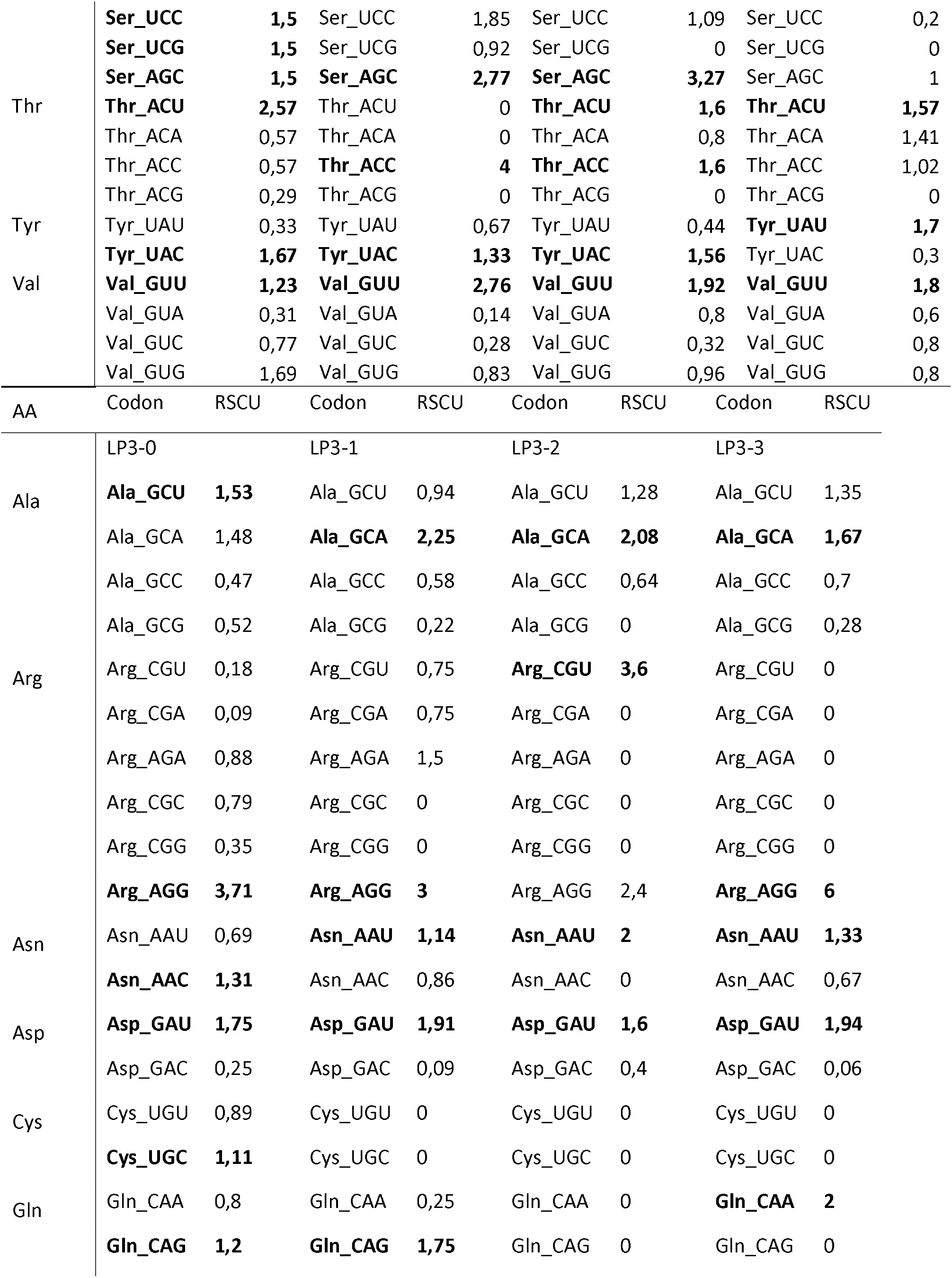
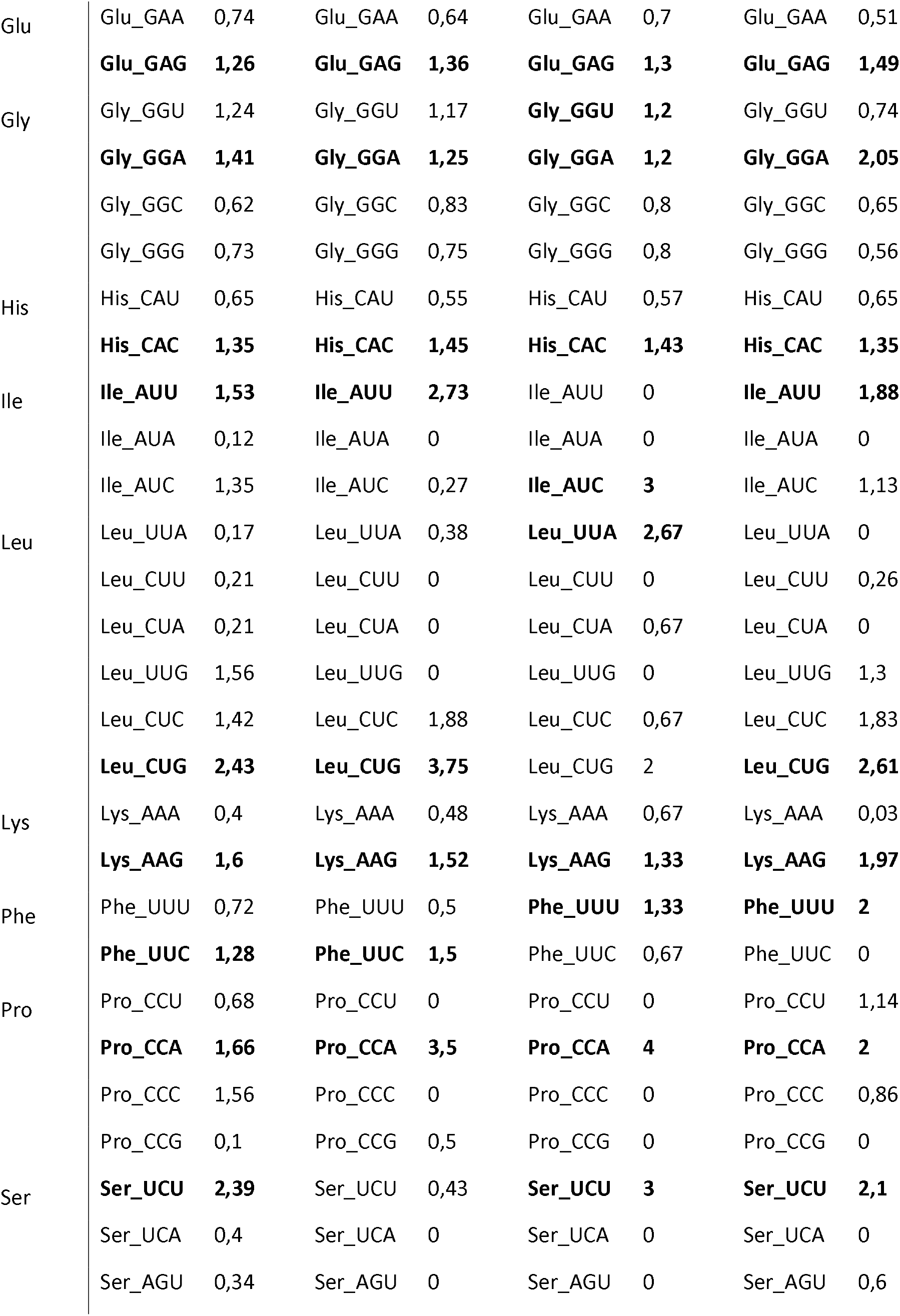
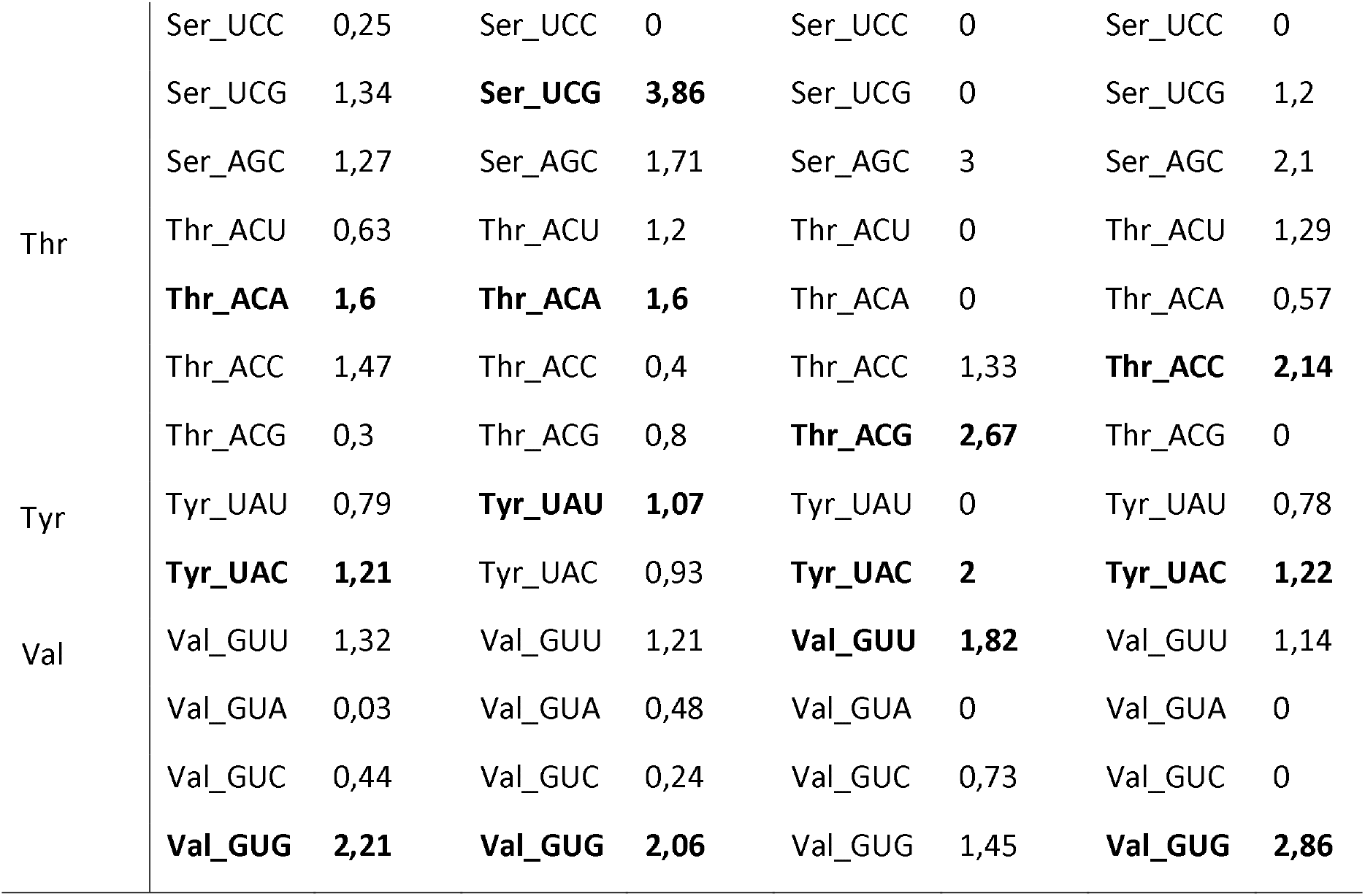
Average RSCU value of each codon per ABA/WDS gene. The most used codon for a given amino acid is indicated in bold.

(a) LP3-0

**Supplementary Table 4:**
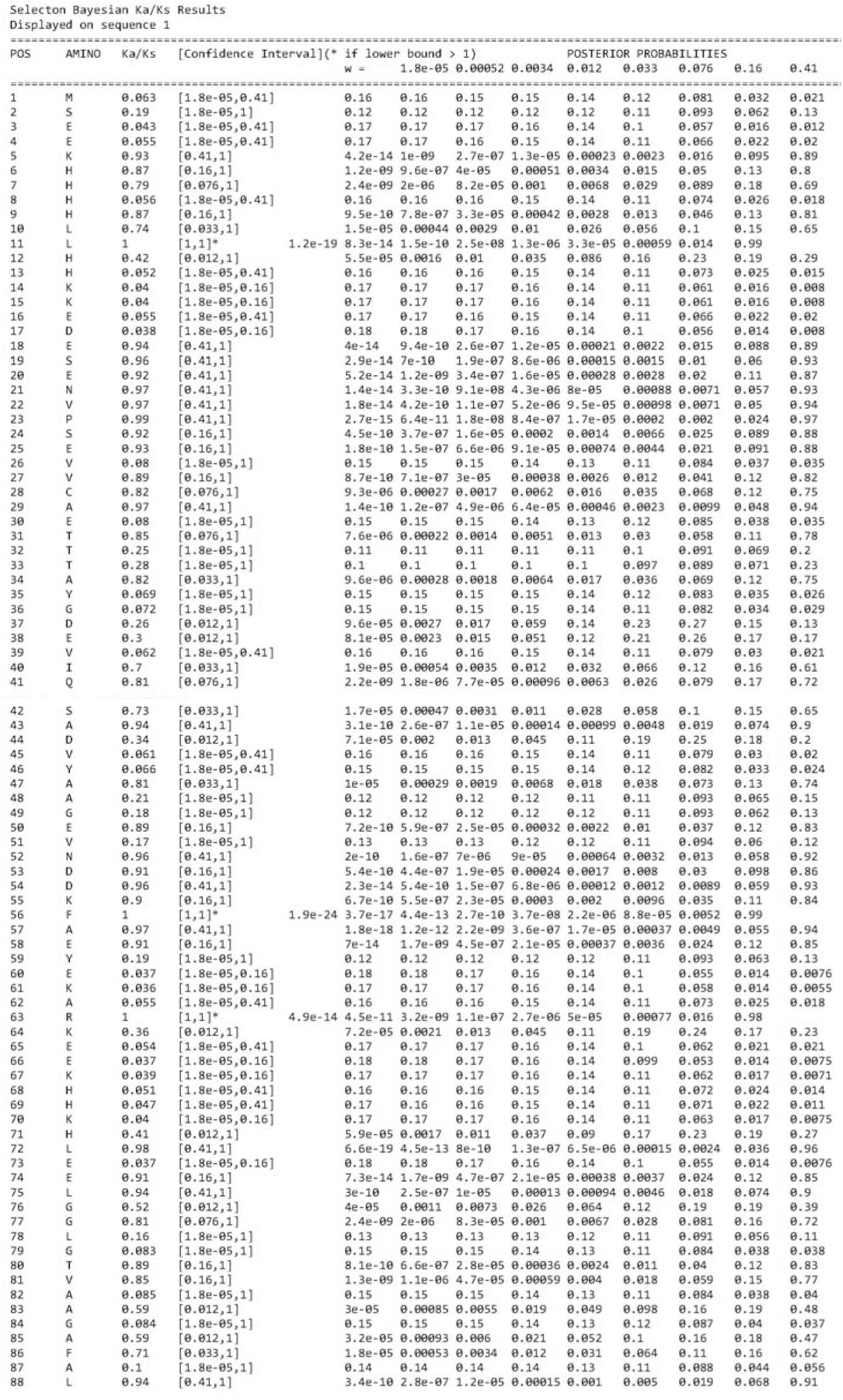
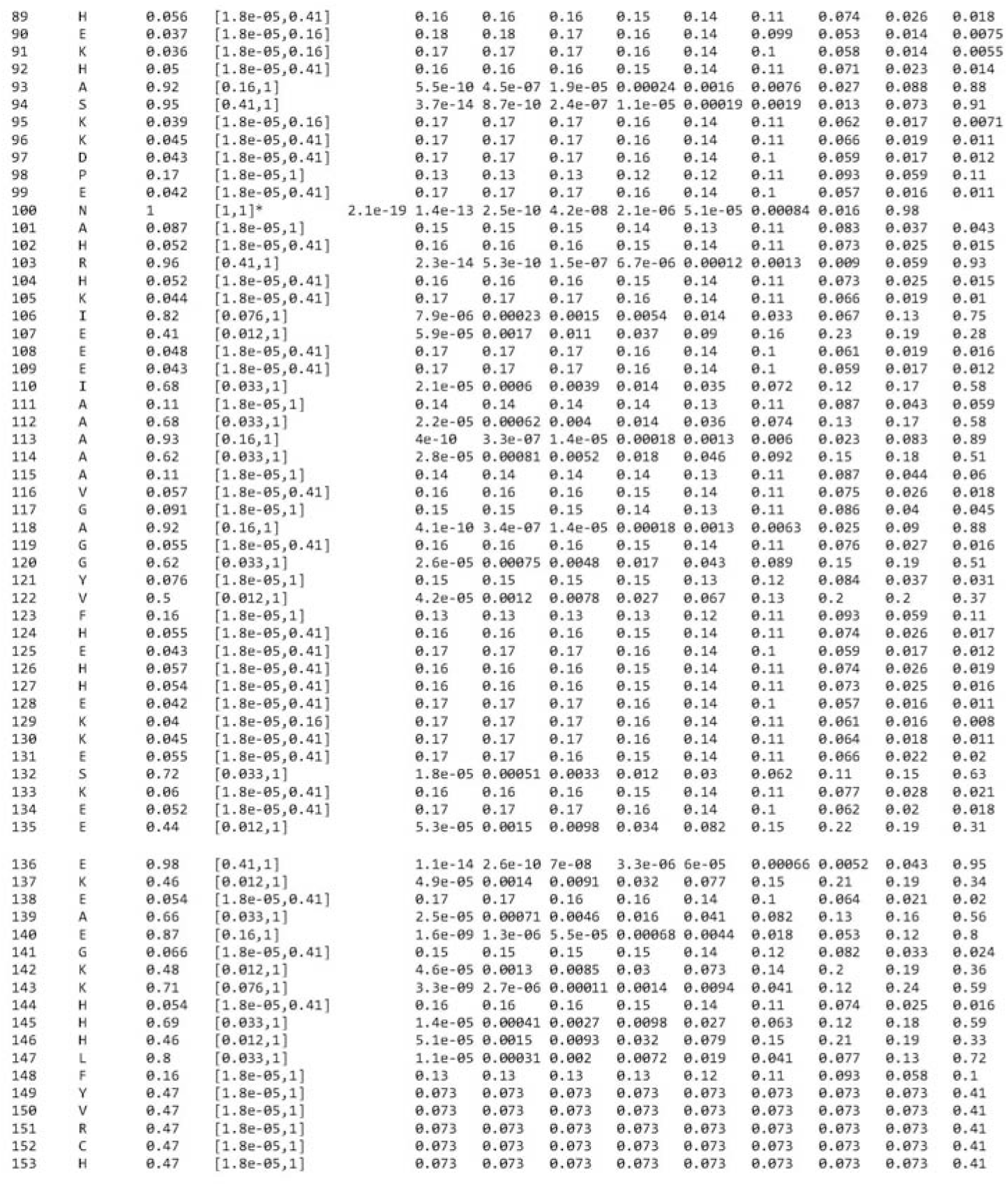
Ka/Ks values per LP3-0 amino acid. First are represented amino acid position, then amino acid, w-score, confidence interval and Bayesian posterior probabilities.

(b) LP3-1

**Supplementary Table 5:**
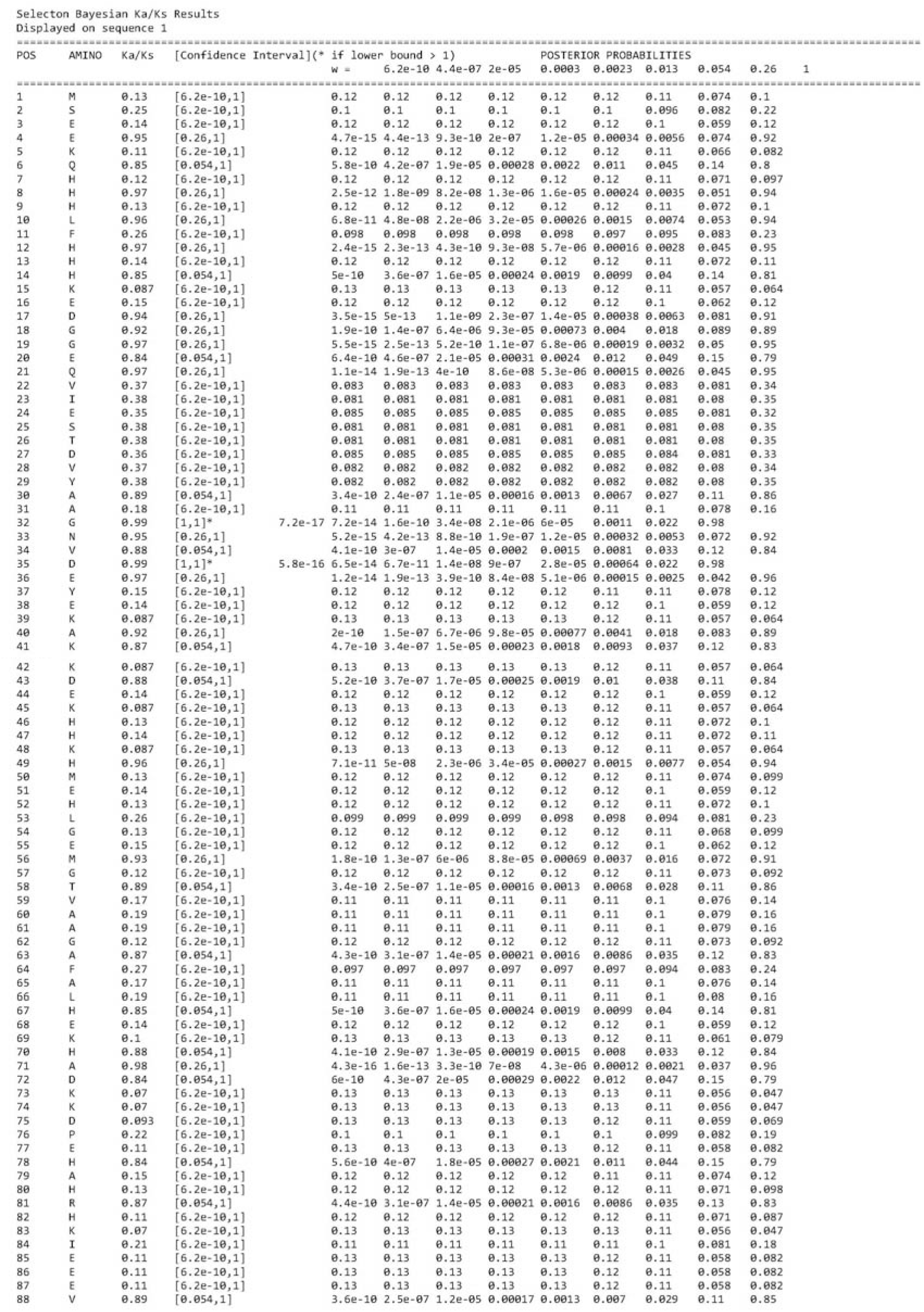
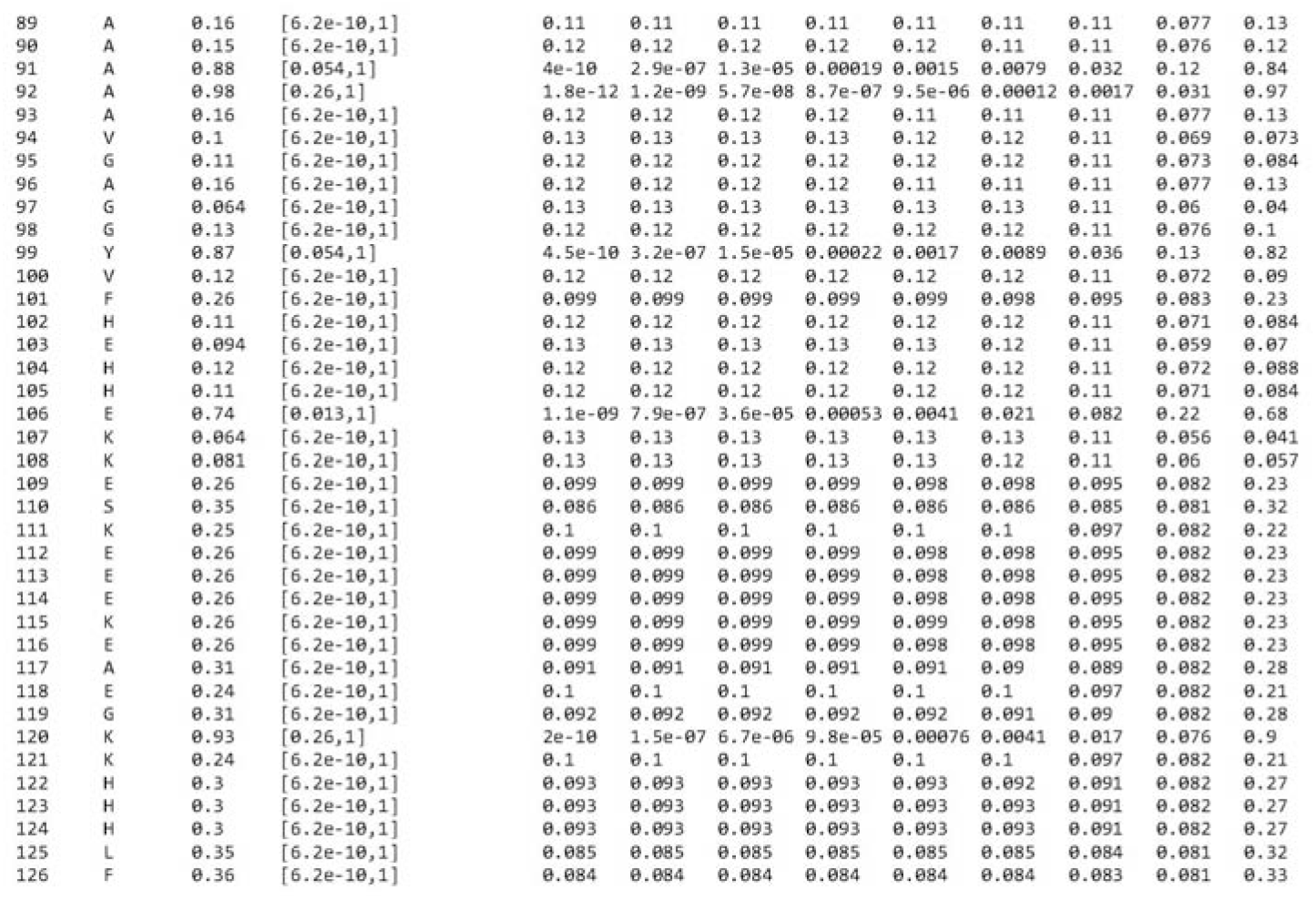
Ka/Ks values per LP3-1 amino acid. First are represented amino acid position, then amino acid, w-score, confidence interval and Bayesian posterior probabilities.

(c) LP3-3

**Supplementary Table 6:**
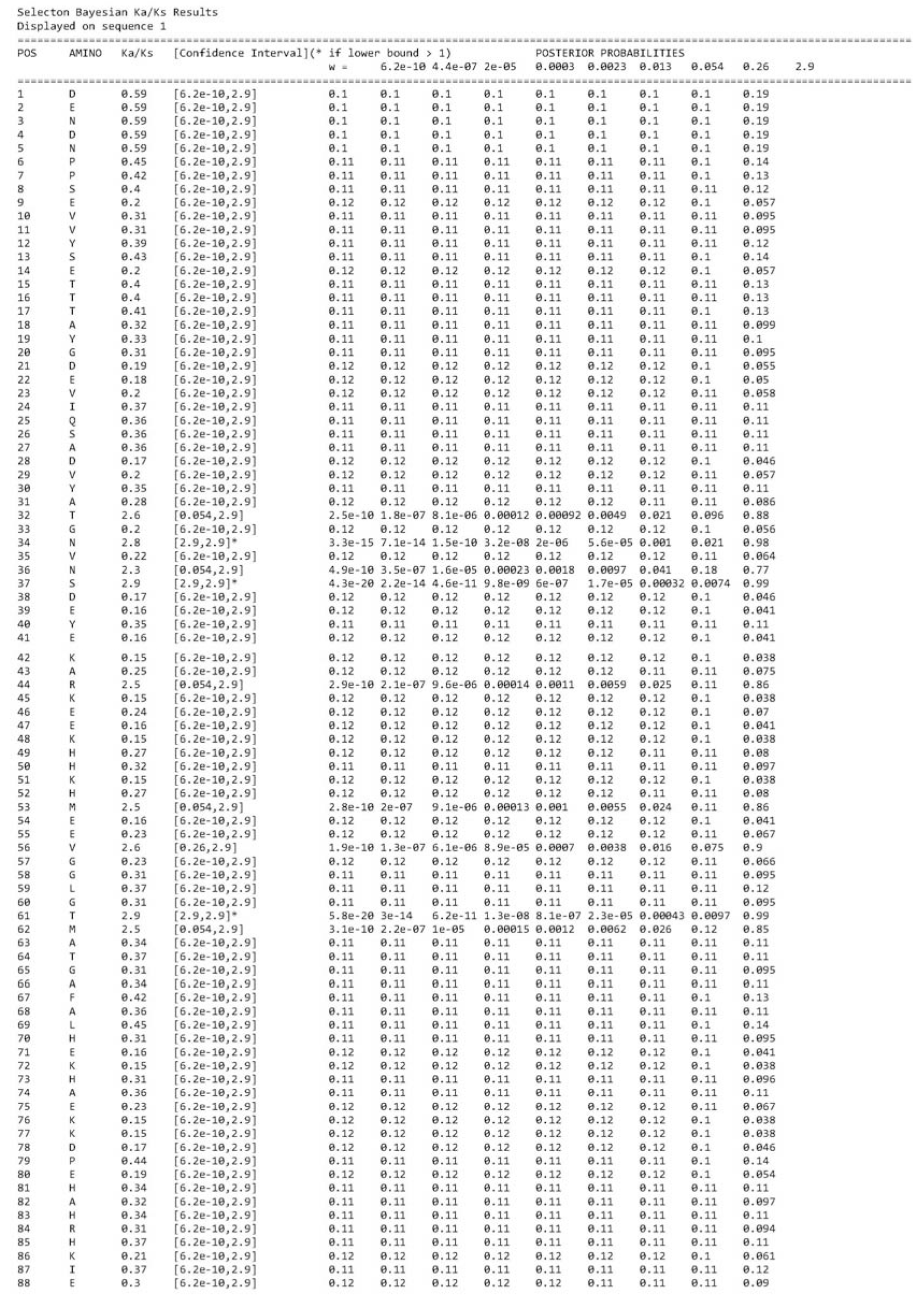
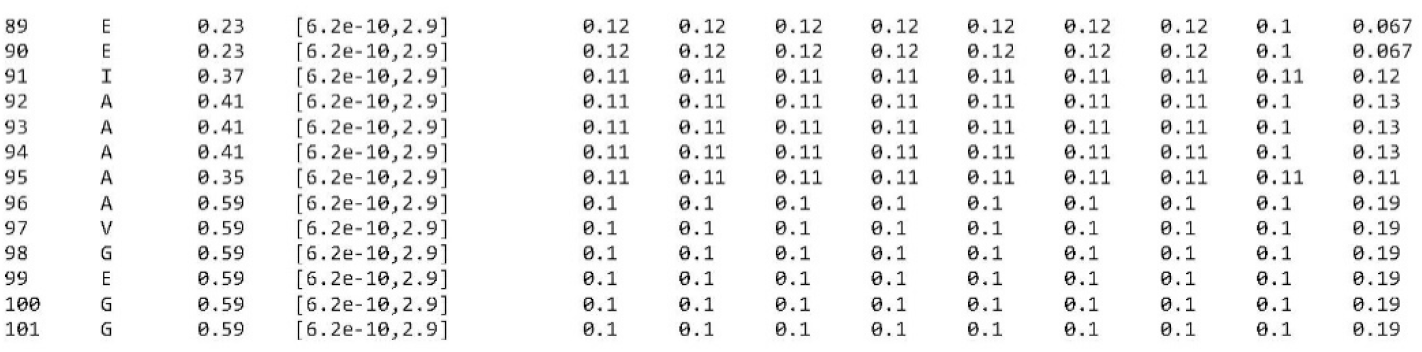
Ka/Ks values per LP3-3 amino acid. First are represented amino acid position, then amino acid, w-score, confidence interval and Bayesian posterior probabilities.

(a) ASR1

**Supplementary Table 7:**
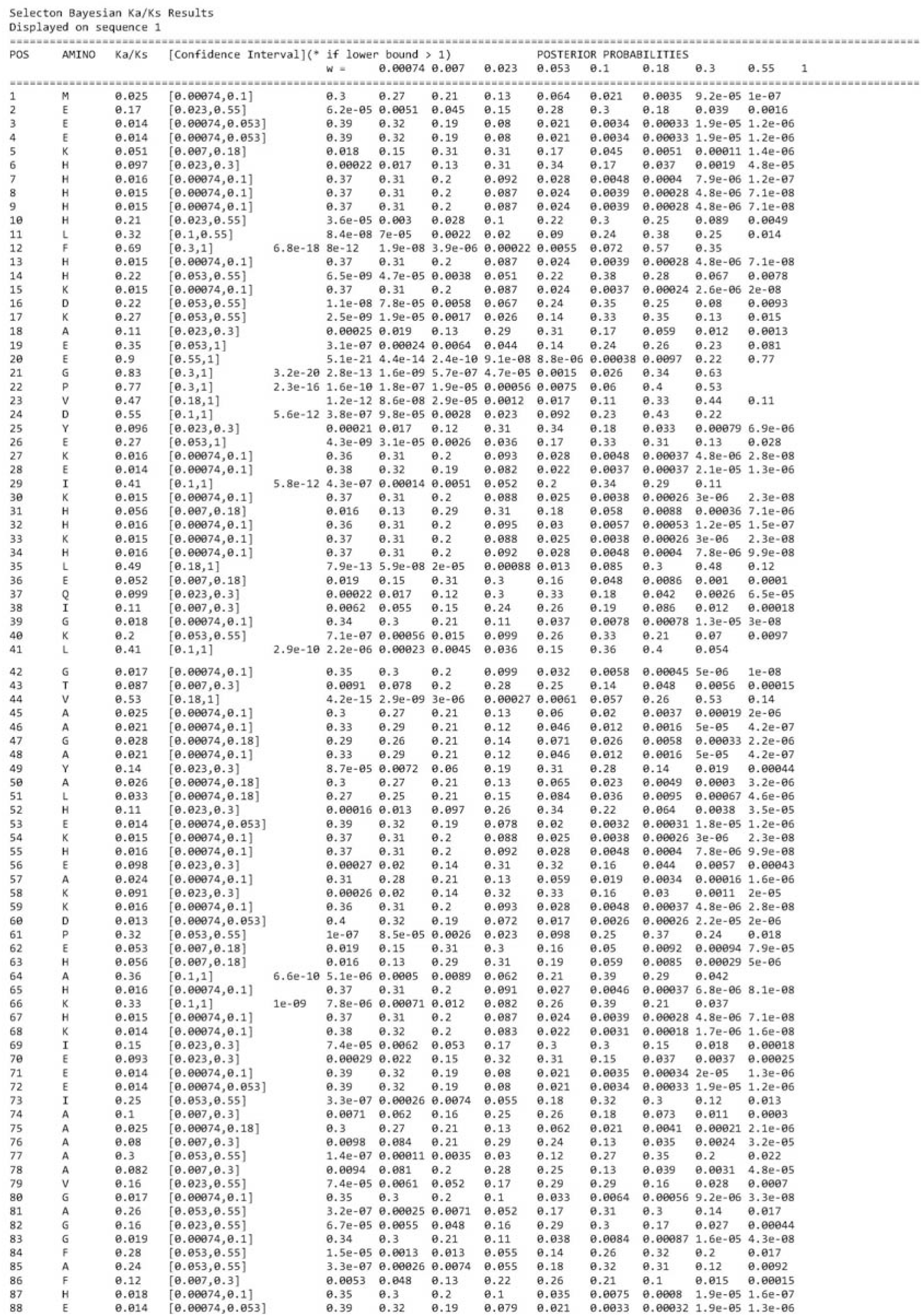
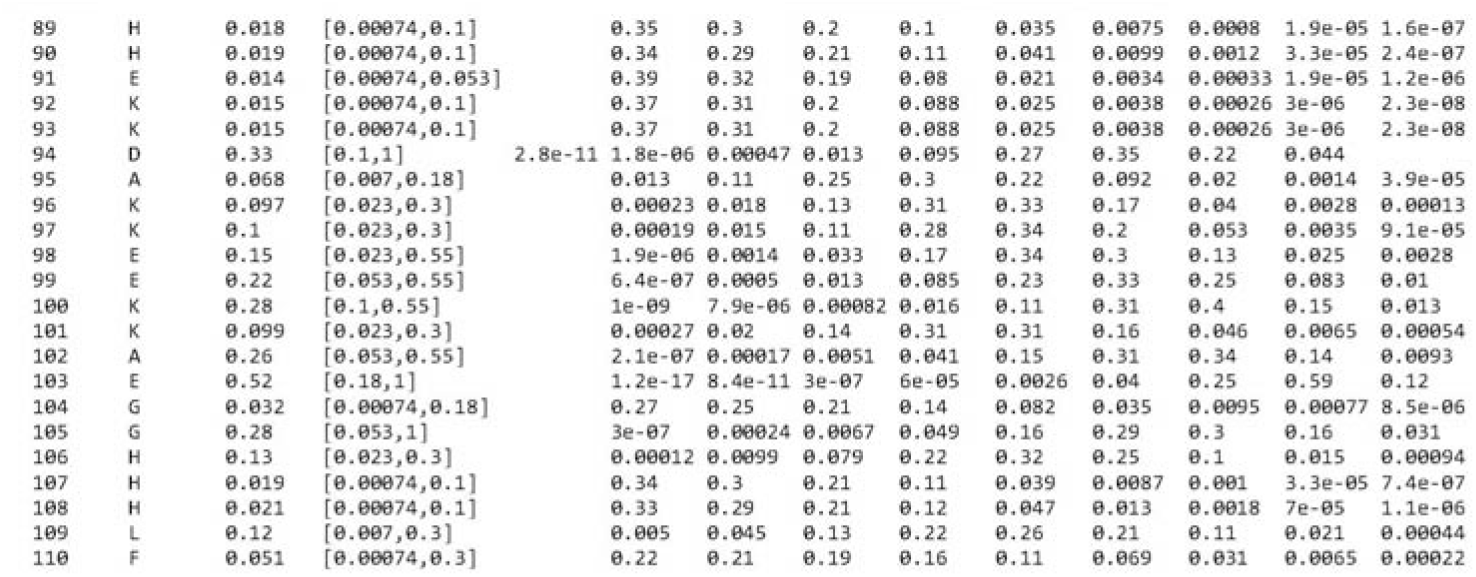
Ka/Ks values per ASR1 amino acid. First are represented amino acid position, then amino acid, w-score, confidence interval and Bayesian posterior probabilities.

(b) ASR2

**Supplementary Table 8:**
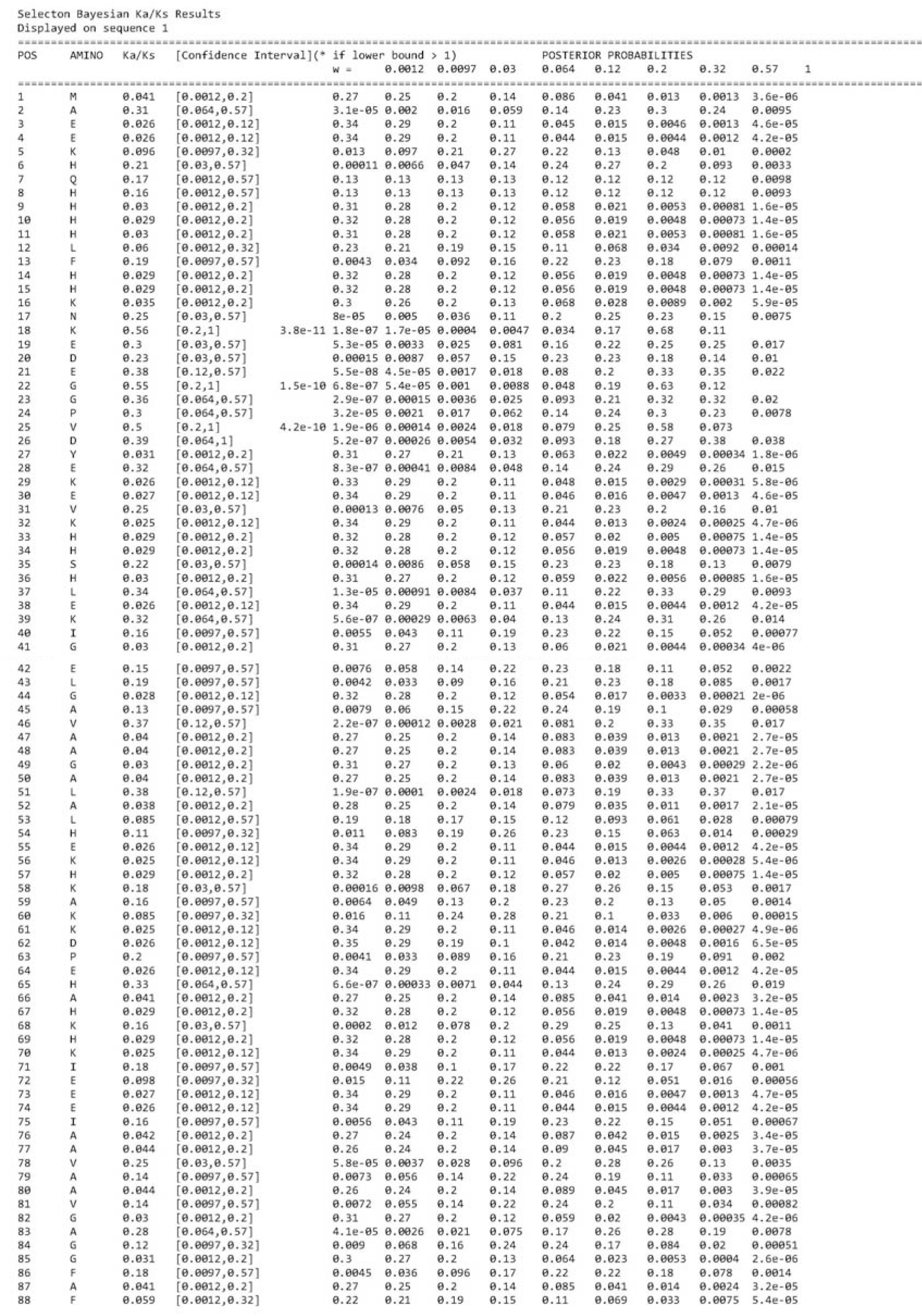
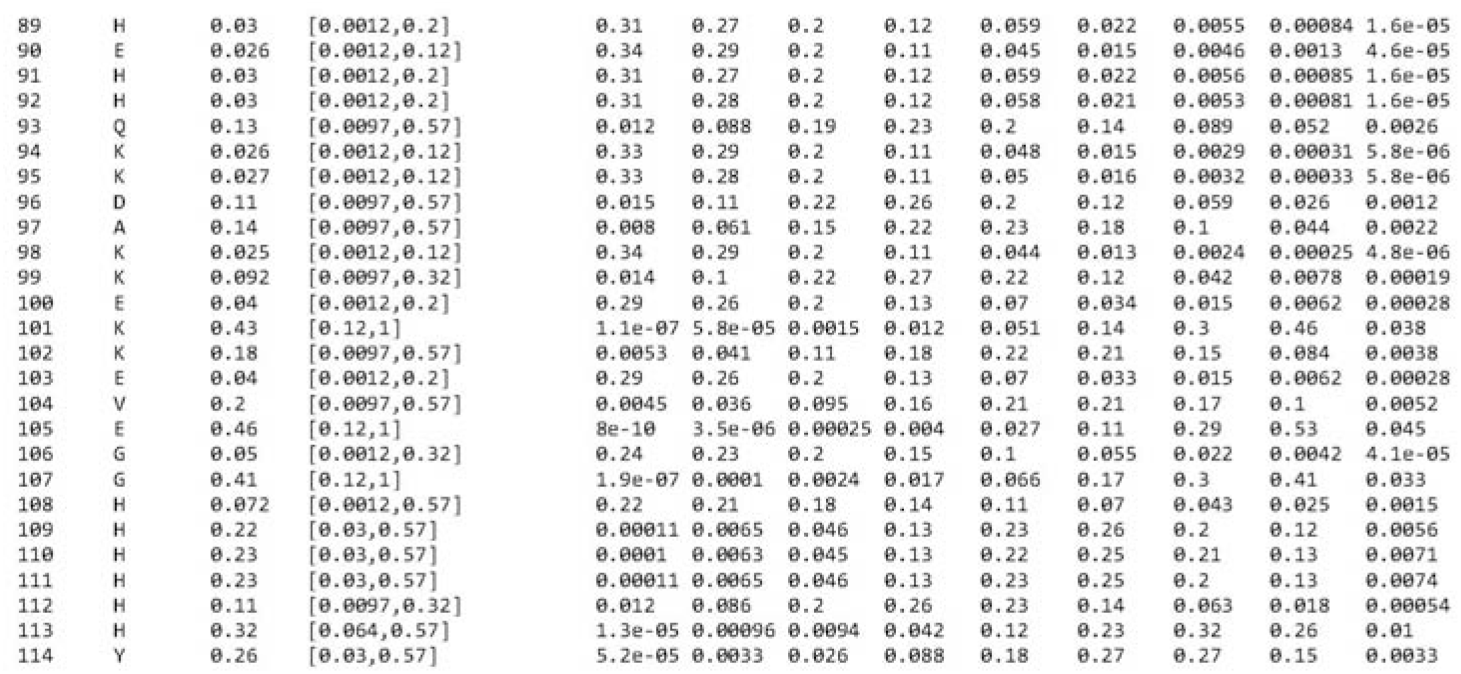
Ka/Ks values per ASR2 amino acid. First are represented amino acid position, then amino acid, w-score, confidence interval and Bayesian posterior probabilities.

(c) ASR3

**Supplementary Table 9:**
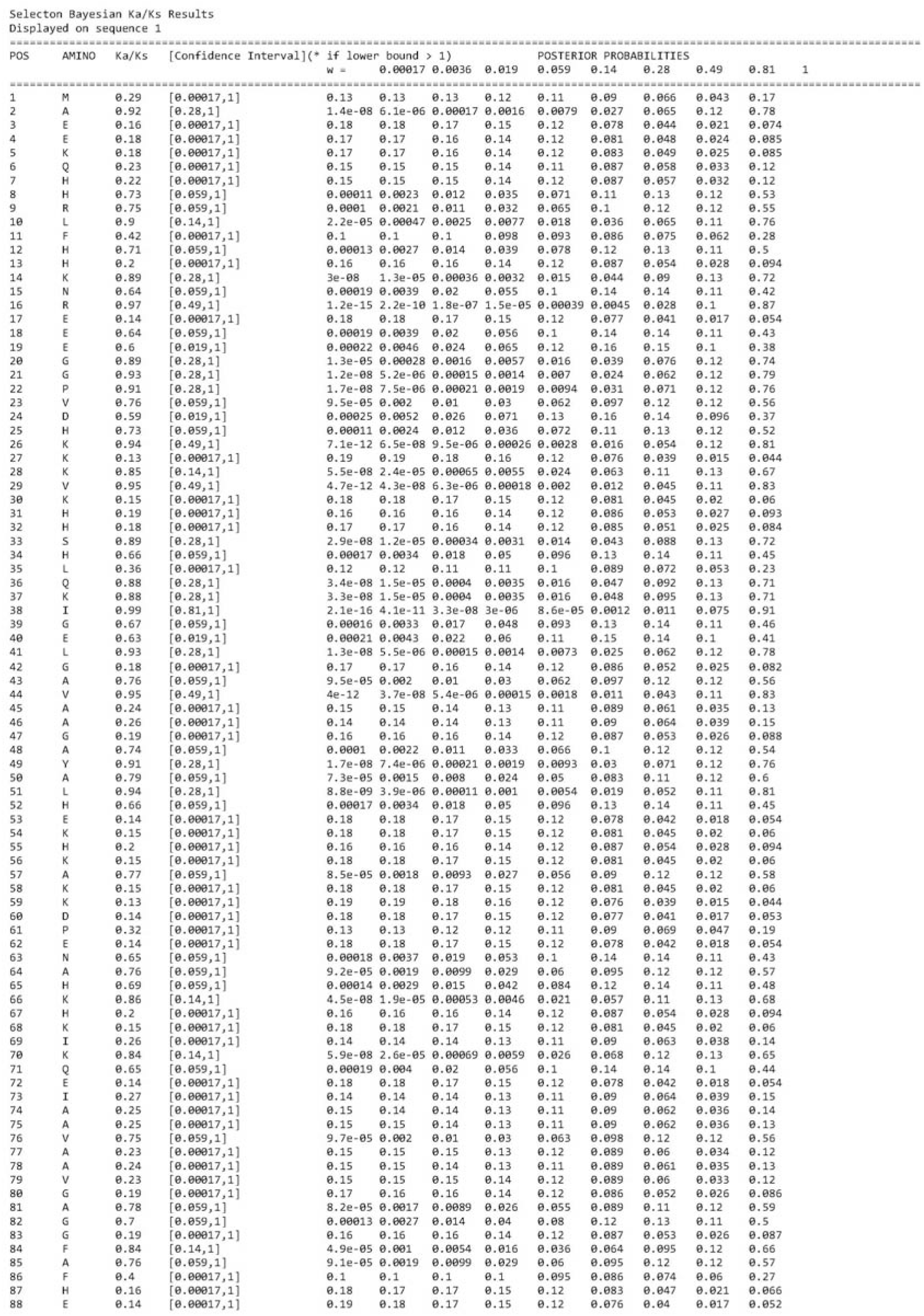
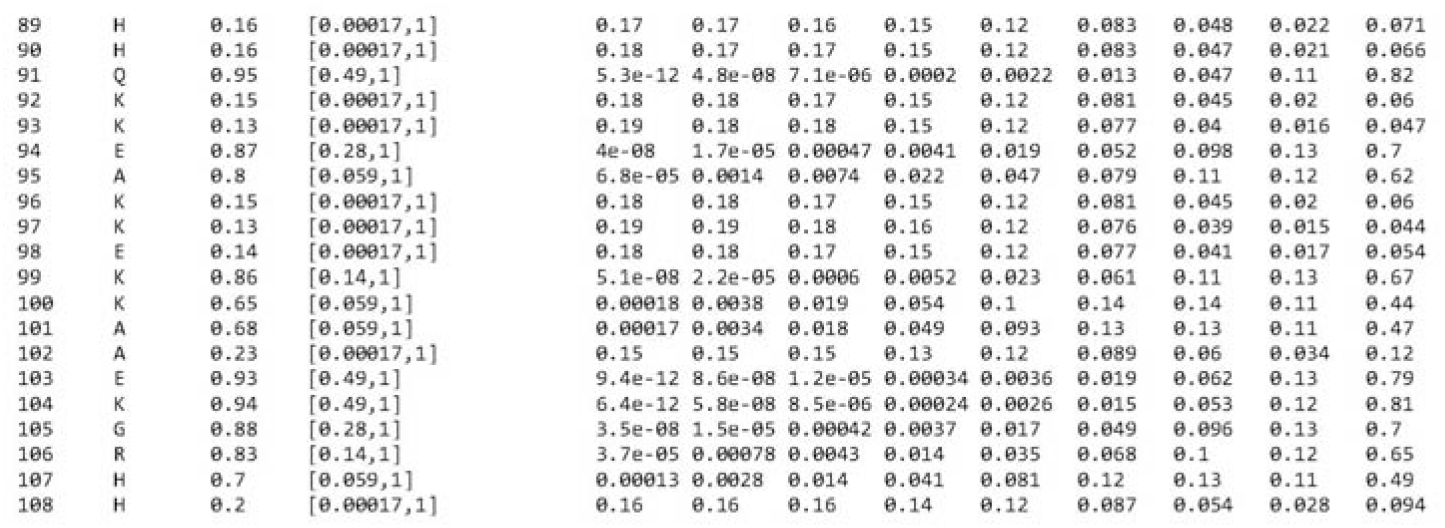
Ka/Ks values per ASR3 amino acid. First are represented amino acid position, then amino acid, w-score, confidence interval and Bayesian posterior probabilities.

(d) ASR4

**Supplementary Table 10:**
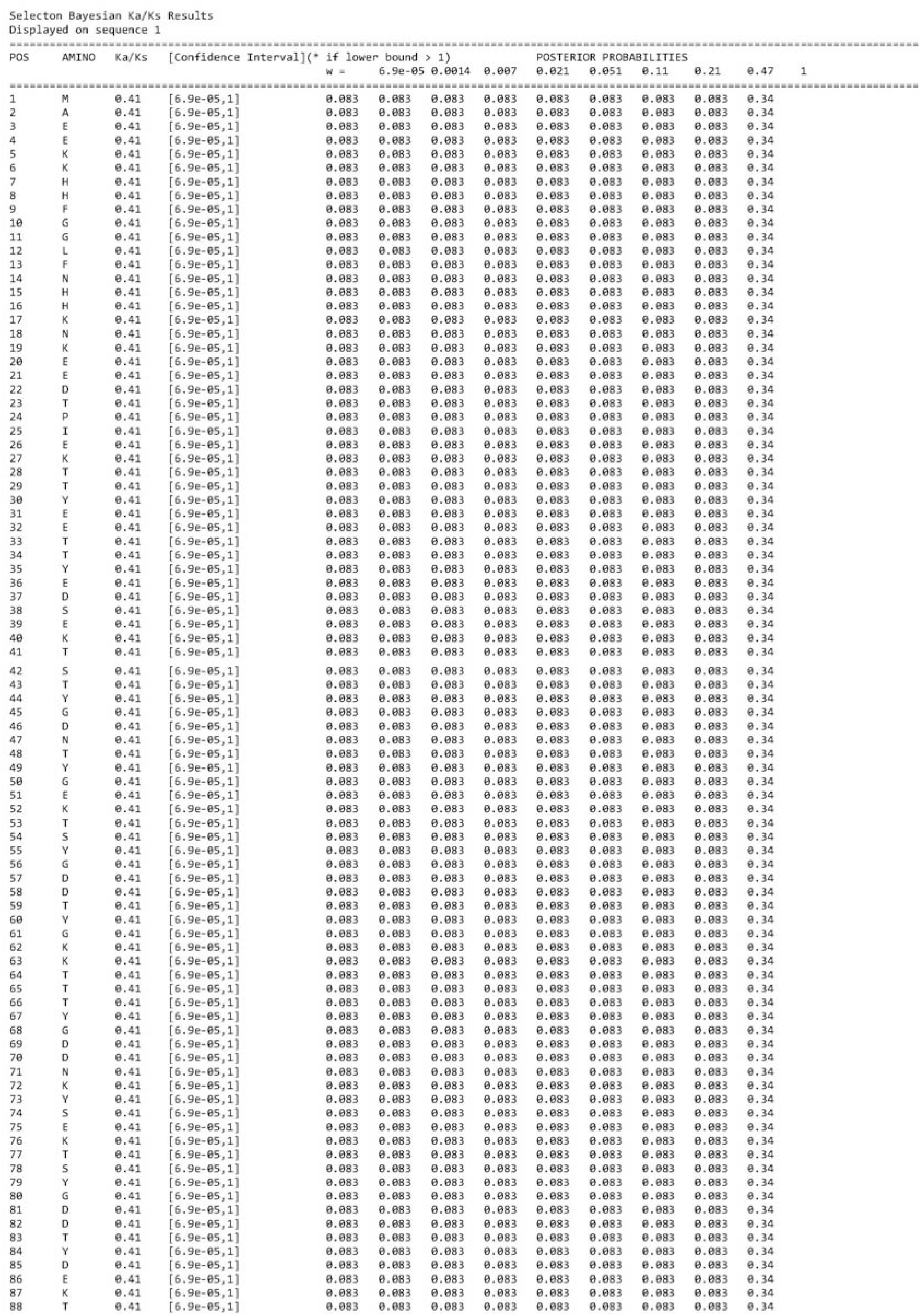
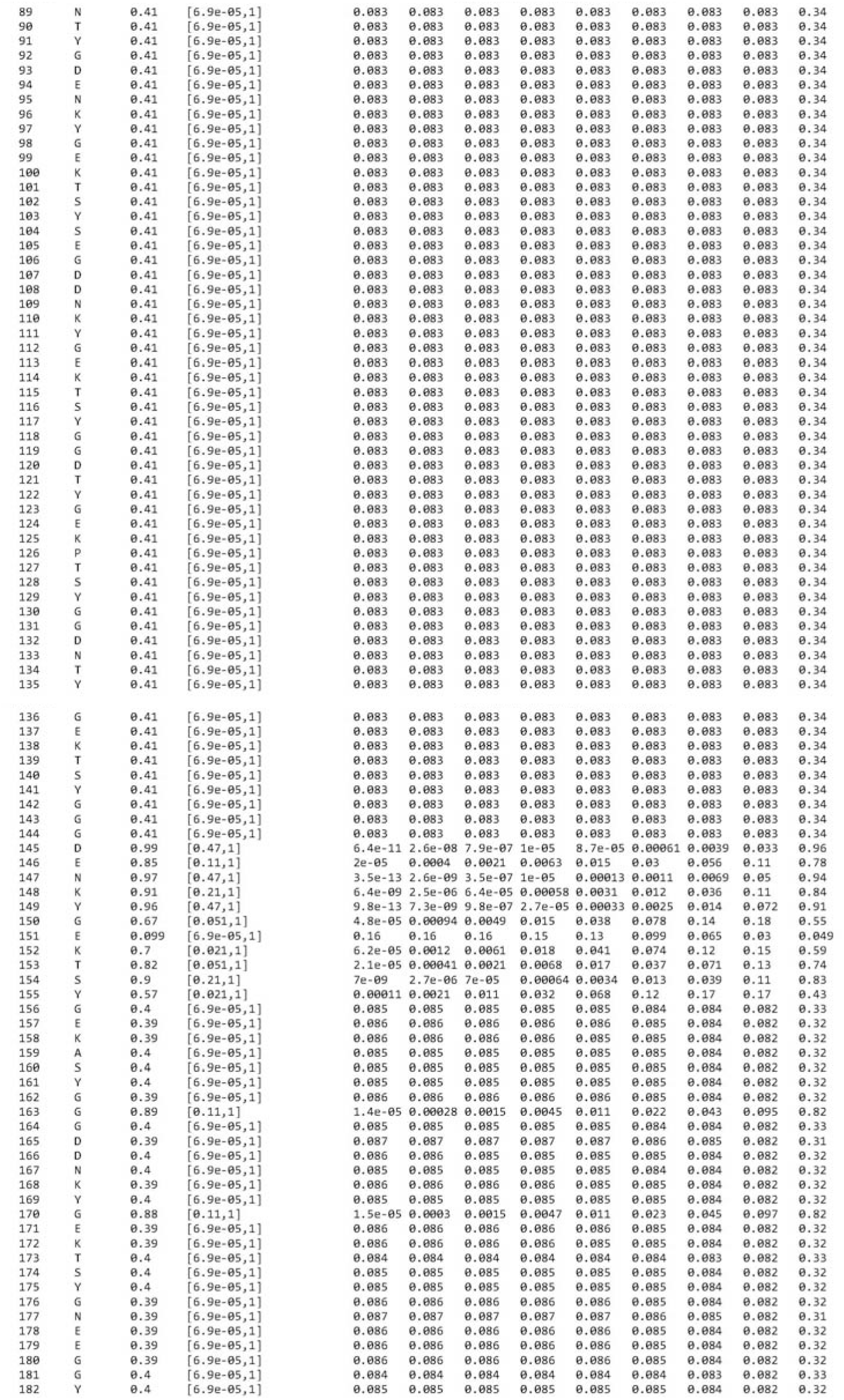
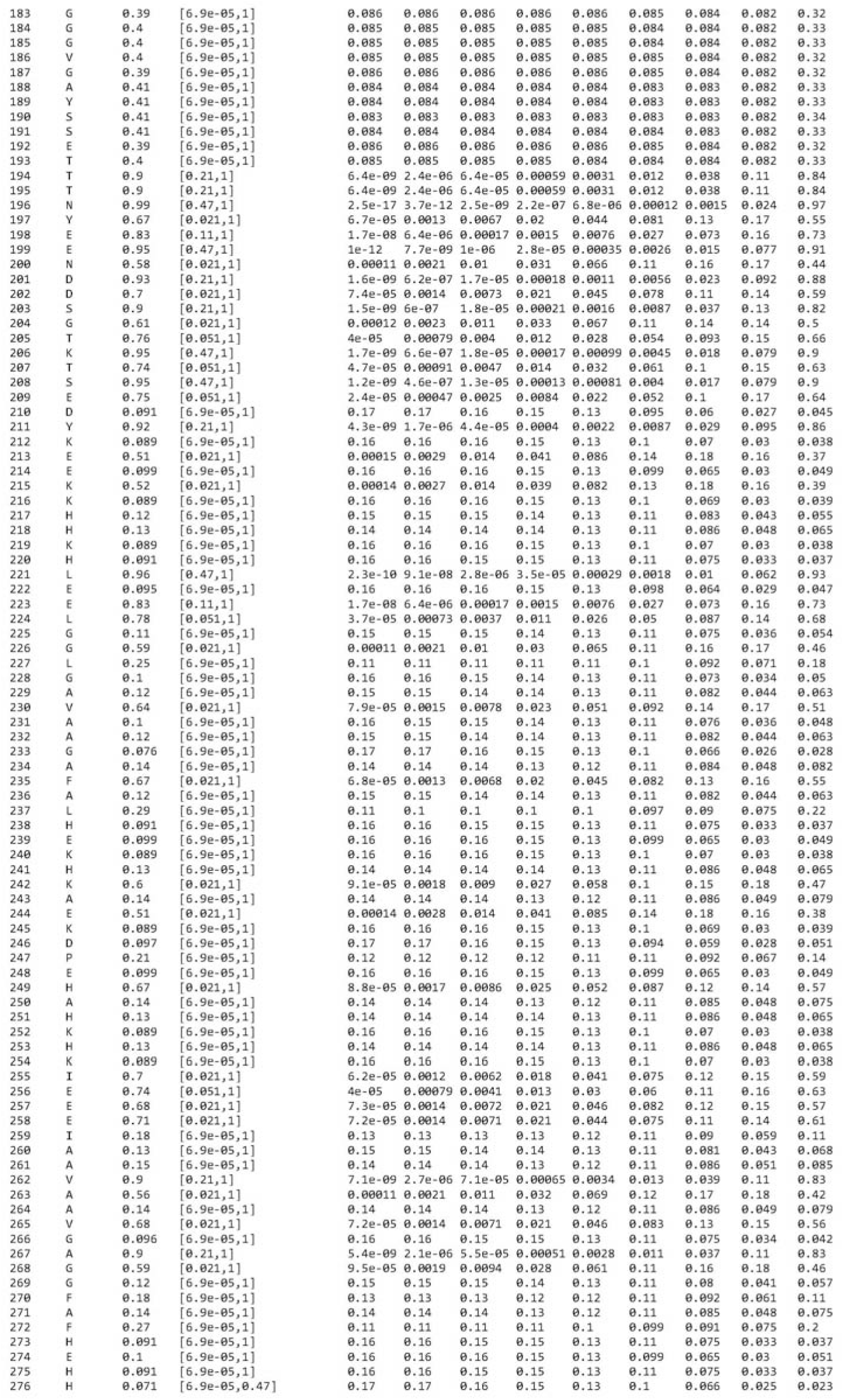
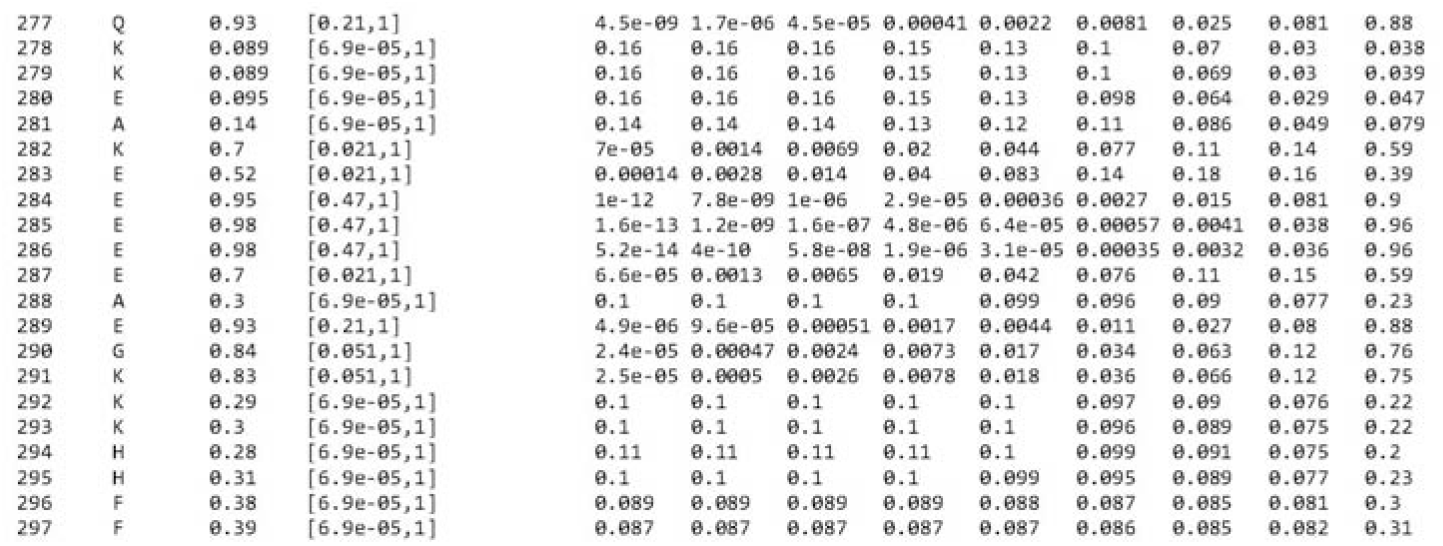
Ka/Ks values per ASR4 amino acid. First are represented amino acid position, then amino acid, w-score, confidence interval and Bayesian posterior probabilities.

