## Supplementary Figures for "Evolutionary analysis of LP3 gene family in conifers: an ASR homolog"

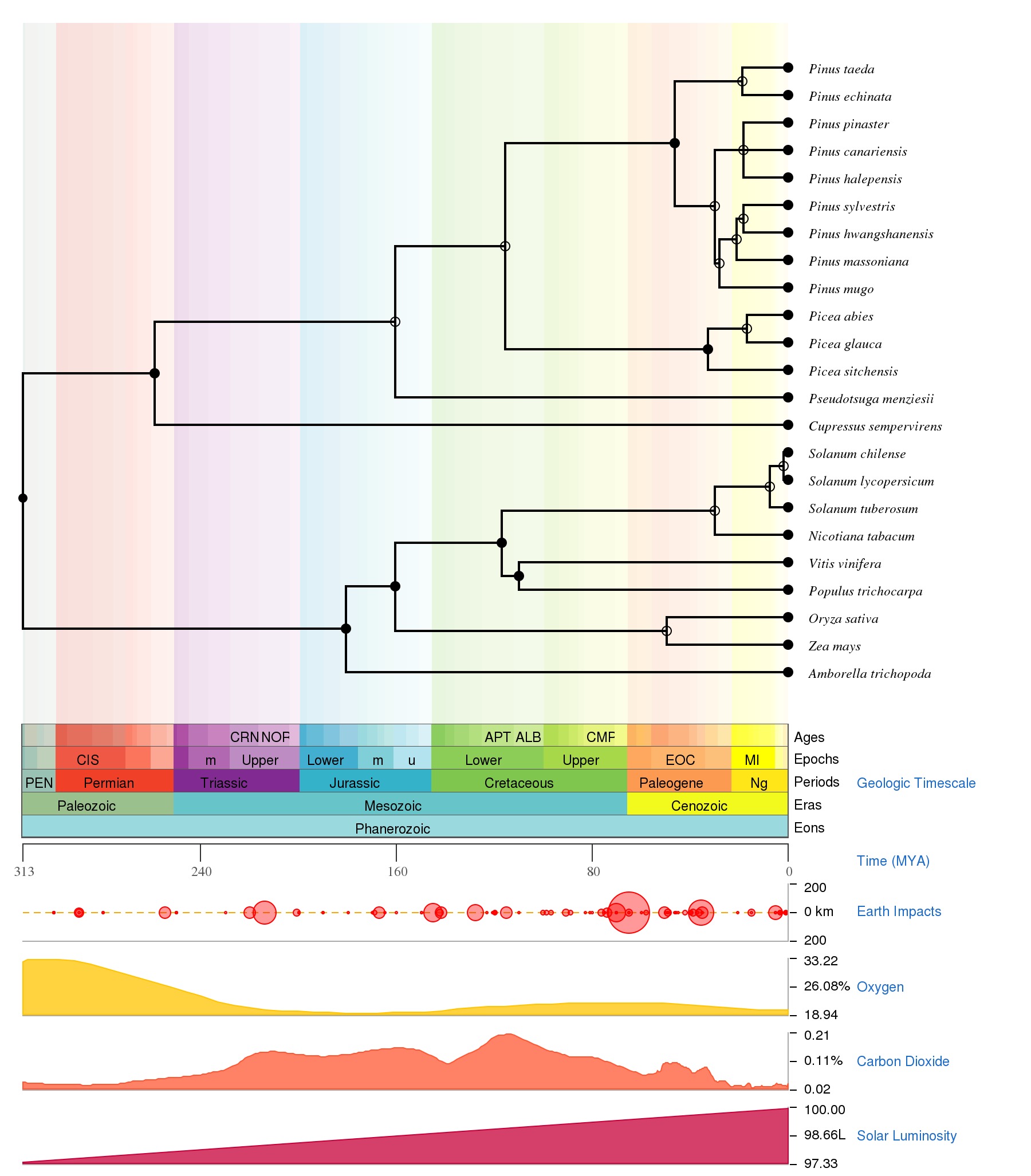

Supplementary Figure 1: Taxonomic phylogenetic tree showing the relationship between the species used in this study. Divergence times are estimated in MYA. Also shown are a geological timescale, meteorite impact magnitude, O2 and CO2 levels in atmosphere and solar luminosity.
